# Mps1 releases Mad1 from nuclear pores to ensure a robust mitotic checkpoint and accurate chromosome segregation

**DOI:** 10.1101/659938

**Authors:** Mariana Osswald, Sofia Cunha-Silva, Jana Goemann, João Barbosa, Luis M Santos, Pedro Resende, Tanja Bange, Claudio E Sunkel, Carlos Conde

## Abstract

The strength of the Spindle Assembly Checkpoint (SAC) depends on the amount of the Mad1-C-Mad2 heterotetramer at kinetochores but also on its binding to Megator/Tpr at nuclear pore complexes (NPCs) during interphase. However, the molecular underpinnings controlling the spatiotemporal redistribution of Mad1-C-Mad2 as cells progress into mitosis remain elusive. Here, we show that Mps1-mediated phosphorylation of Megator/Tpr abolishes its interaction with Mad1 *in vitro* and in *Drosophila* cells. Timely activation of Mps1 during prophase triggers Mad1 release from NPCs, which we find to be required for competent kinetochore recruitment of Mad1-C-Mad2 and robust checkpoint response. Importantly, preventing Mad1 binding to Megator/Tpr rescues the fidelity of chromosome segregation and aneuploidy in larval neuroblasts of *Drosophila mps1*-null mutants. Our findings demonstrate that the subcellular localization of Mad1 is stringently coordinated with cell cycle progression by kinetochore-extrinsic activity of Mps1. This ensures that both NPCs in interphase and kinetochores in mitosis can generate anaphase inhibitors to efficiently preserve genomic stability.

## INTRODUCTION

The Spindle Assembly Checkpoint (SAC) safeguards eukaryotic cells against chromosome mis-segregation by restraining the transition to anaphase in the presence of unattached kinetochores. Pivotal to this signalling pathway, is the Mad1-Mad2 heterotetramer that catalyses the structural conversion of soluble open-Mad2 (O-Mad2) into closed-Mad2 (C-Mad2), a conformer that is able to bind the APC/C activator Cdc20 (De Antoni et al., 2005). This represents the rate-limiting step in the assembly of the mitotic checkpoint complex (MCC), a diffusible tetrameric complex that inhibits APC/C-mediated ubiquitination of securin and cyclin B and thereby delays sister chromatid separation and mitotic exit (De Antoni et al., 2005; Simonetta et al., 2009; Faesen et al., 2017). Compelling evidence indicate that the strength of the SAC response is dictated by the amount of Mad1-C-Mad2 present at kinetochores (Collin et al., 2013; Dick and Gerlich, 2013; Hustedt et al., 2014). However, a sustained SAC signalling also requires Mad1-C-Mad2 to associate with nuclear pore complexes (NPCs) during interphase, which is mediated through Mad1 binding to the nuclear basket nucleoporin Megator/Tpr (Scott et al., 2005; Lee et al., 2008; Souza et al., 2009; Lince-Faria et al., 2009; Schweizer et al., 2013; Rodriguez-Bravo et al., 2014). This arrangement regulates Mad1-C-Mad2 proteostasis to ensure that sufficient amount of complexes are produced before mitosis (Schweizer et al., 2013). Moreover, it was proposed that Mad1-C-Mad2 at NPCs also activates O-Mad2 into C-Mad2, hence providing a scaffold for the assembly of pre-mitotic MCC (Rodriguez-Bravo et al., 2014). This is thought to operate as a mitotic timer to support APC/C inhibition during early mitosis until newly formed kinetochores are able to instate efficient SAC activation (Sudakin et al., 2001; Meraldi et al., 2004; Malureanu et al., 2009; Maciejowski et al., 2010; Rodriguez-Bravo et al., 2014; Kim et al., 2018). Notwithstanding its importance for mitotic fidelity, how the subcellular redistribution of Mad1-C-Mad2 is coordinated with cell cycle progression remains elusive. Particularly, whether and how regulatory events at NPCs impact on Mad1-C-Mad2 kinetochore localization has not been established so far. We set out to address these questions in *Drosophila*, where the multi-sequential phosphorylation cascade controlling Mad1 kinetochore localization through the Mps1-Knl1-Bub1 pathway (London et al., 2012; Shepperd et al., 2012; Yamagishi et al., 2012; Primorac et al., 2013; London and Biggins, 2014; Vleugel et al., 2015; Mora-Santos et al., 2016; Faesen et al., 2017; Ji et al., 2017; Qian et al., 2017; Zhang et al., 2017; Rodriguez-Rodriguez et al., 2018) is inherently absent (Schittenhelm et al., 2009; Conde et al., 2013). This reduces kinetochore-associated complexity, hence providing a simpler naturally occurring system to uncover the potential role of kinetochore-extrinsic mechanisms in Mad1-C-Mad2 subcellular distribution throughout the cell-cycle and their significance for SAC signalling and genomic integrity *in vivo.*

## RESULTS AND DISCUSSION

### Mps1 triggers Mad1 exclusion from nuclear pore complexes during prophase

To investigate the events underlying the subcellular redistribution of Mad1 during mitotic entry we first monitored with high-temporal resolution the dynamics of Mad1 and Megator localization in *Drosophila* S2 cells (Figure 1A,B and S1A,B). Mad1-EGFP signal at the nuclear envelope (NE) begins to fade during early prophase whereas Megator-EGFP intensity persists until tubulin becomes detectable in the nucleus, an early event of nuclear envelope breakdown (NEB). Interestingly, depletion of Mps1 causes a significantly delay in Mad1-EGFP dissociation from the NE with no discernible impact on Megator-EGFP dynamics (Figure 1A,B and S1A,B). The decline in Mad1-EGFP signal intensity at the NE of Mps1-depleted cells entering mitosis overlaps perfectly with the pattern of Megator-EGFP, hinting that in the absence of Mps1 activity, the exclusion of Mad1 from NPCs is restrained by the presence of the nucleoporin (Figure S1A,B). These results support that Mad1 reallocation from NPCs is triggered before NEB onset in an Mps1-dependent manner. Consistently, a phospho-specific antibody recognizing the activating autophosphorylation (T490Ph) of Mps1 T-loop (Jelluma et al., 2008; Moura et al., 2017) decorates the NE throughout prophase, thus indicating that Mps1 is active at NPCs during mitotic entry. We then tested whether inducing Mps1 activation in interphase cells prematurely displaces Mad1 from NPCs. As Mps1 is excluded from the nucleus until late G2/early prophase (Zhang et al., 2011; Jia et al., 2015), we promoted its nuclear import by fusing it with the SV40 large T-antigen nuclear localization signal (EGFP-Mps1^WT^-NLS). Strikingly, overexpression of EGFP-Mps1^WT^-NLS efficiently elicits nuclear activation of Mps1 (Figure S1C) and clearly decreases Mad1 levels at NPCs of interphase cells (Figure 1D,E). In contrast, Mad1 association with NPCs remains unaltered in interphase S2 cells overexpressing catalytic dead EGFP-Mps1^KD^-NLS or EGFP-Mps1^WT^ (Figure 1D,E), which albeit active in the cytoplasm, fails to attain close proximity with the nucleoplasmic side of NPCs (Figure S1C). Collectively, these results demonstrate that timely control of Mps1 nuclear import and activation triggers Mad1 dissociation from NPCs during early prophase before NEB onset.

**Figure 1:**
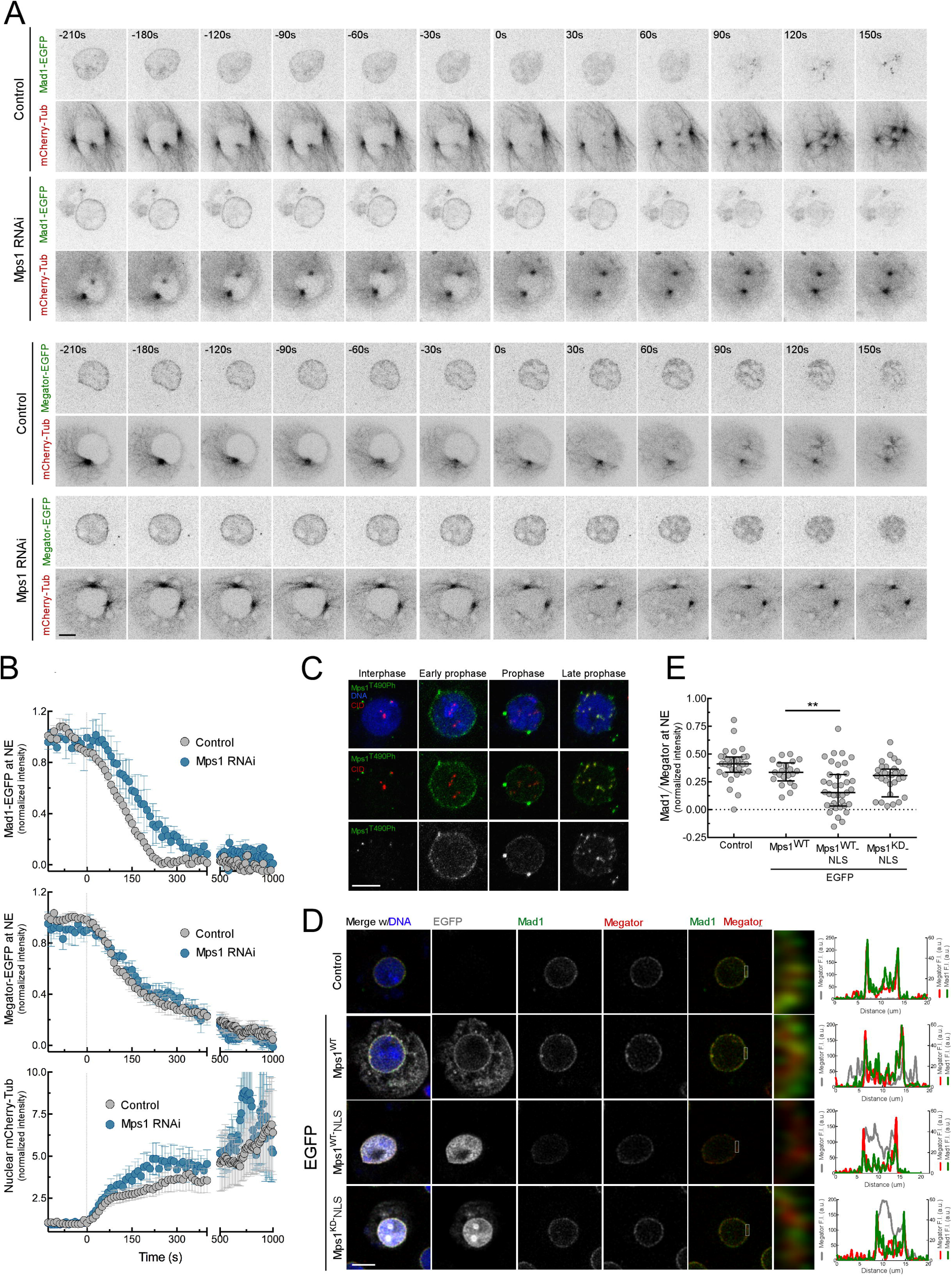
Msp1 promotes the dissociation of Mad1 from nuclear pore complexes during prophase. (A,B) Representative mitotic progression (A) and quantification (B) of Mad1-EGFP or Megator-EGFP levels at nuclear envelope (NE) and of mCherry-Tubulin levels in the nucleus of control and Mps1-depleted *Drosophila* S2 cells. Mitotic progression was monitored through time-lapse microscopy. Time 0s indicates nuclear envelope breakdown (NEB) and was defined as the moment mCherry-Tubulin signal becomes detectable in the nucleus. Mad1-EGFP (N ≥7 cells), Megator-EGFP (N ≥6 cells) and mCherry-Tubulin fluorescence intensities were normalized to the mean value before NEB. (C) Representative immunofluorescence images of Mps1^T490Ph^ localization pattern in interphase and prophase S2 cells. (D, E) Representative immunofluorescence images (D) and quantifications (E) of Mad1 levels at the nuclear envelope of interphase control S2 cells and interphase S2 cells expressing EGFP-Mps1^WT^, EGFP-Mps1^WT^-NLS or EGFP-Mps1^KD^-NLS. The insets display magnifications of the outlined regions. Mad1 fluorescence intensities at the nuclear envelope were determined relative to Megator signal (N ≥ 21 cells for each condition). Data information: in (B) data is presented as mean ± SD, in (E) data is presented as median with the interquartile range. Asterisks indicate that differences between mean ranks are statistically significant, **p<0.005 (Kruskal-Wallis, Dunn’s multiple comparison test). Scale bars: 5μm.

### Mps1-mediatedphosphorylation of Megator disrupts its interaction with Mad1

Since Mad1 localization at NPCs is mediated by Megator (Scott et al., 2005; Lee et al., 2008; Souza et al., 2009; Lince-Faria et al., 2009; Schweizer et al., 2013; Rodriguez-Bravo et al., 2014) we next sought to examine whether Mps1 activity directly affects this interaction. We found that Megator preferentially co-immunoprecipitates with Mad1 when Mps1 is depleted from mitotically-enriched S2 cells lysates (Figure 2A). Moreover, knocking-down the kinase markedly reduced Megator hyperphosphorylation (Figure 2B), which prompt us to examine whether Megator is directly targeted by Mps1. *In vitro* kinase assays and mass-spectrometry analysis using recombinant Mps1 and fragments of Megator N-terminus potentially involved in Mad1 binding (Lee et al., 2008) revealed that GST-Mps1 phosphorylates MBP-Megator^1178-1655^ on T1259, T1295, T1338 and T1390. These residues are located in a putative coiled-coil region (Figure 2D), which we found to directly interact with Mad1 N-terminus in pull-down assays (Figure 2E,F). Notably, 6xHis-Mad1^1-493^ failed to bind with the same efficiency to MBP-Megator^1178-1655/WT^ that had been previously phosphorylated by GST-Mps1 or to a phosphomimetic version where T1259, T1295, T1338 and T1390 are converted to aspartates (MBP-Megator^1178-1655/T4D^), thus indicating that phosphorylation of these particular residues negatively regulates Mad1 binding to Megator *in vitro* (Figure 2E,F). To monitor Mad1-Megator interaction in mitotic cells, we resorted to light-activated reversible inhibition by assembled trap (LARIAT). With this optogenetic tool, EGFP-tagged proteins are sequestered into complexes formed by a multimeric protein (MP) and a blue light-mediated heterodimerization Cib1-Cry2 module (Figure 2G,H). MP is fused to the cryptochrome-interacting basic helix-loop-helix 1 (Cib1), to which the cryptochrome Cry2 is able to bind when photoactivated (Kennedy et al., 2010; Lee et al., 2014; Osswald et al., 2019). By tagging Cry2 with an anti-GFP nanobody, we were able to induce clustering of wild-type (WT), phosphodefective (T4A) and phosphomimetic (T4D) versions of EGFP-Megator^1178-1655^ with high spatiotemporal resolution and examine their capacity to recruit Mad1 (Figure 2G,H). Immunofluorescence analysis reveals limited association of Mad1 with clusters of EGFP-Megator^1178-1655/WT^ present in the cytoplasm of colchicine-treated S2 cells (Figure 2G,I). However, a significant increment in Mad1 recruitment to clustered EGFP-Megator^1178-1655/WT^ occurs upon depletion of Mps1 and similar levels of Mad1 are observed at clusters of EGFP-Megator^1178-1655/T4A^ (Figure 2G,I). Importantly, Mad1 fails to associate with clusters of EGFP-Megator^1178-1655/T4D^, even after Mps1 knock-down (Figure 2G,I). Collectively, these results demonstrate that Mps1-mediated phosphorylation of Megator on T1259, T1395, T1338 and T1390 prevents it from binding to Mad1 during mitosis.

**Figure 2:**
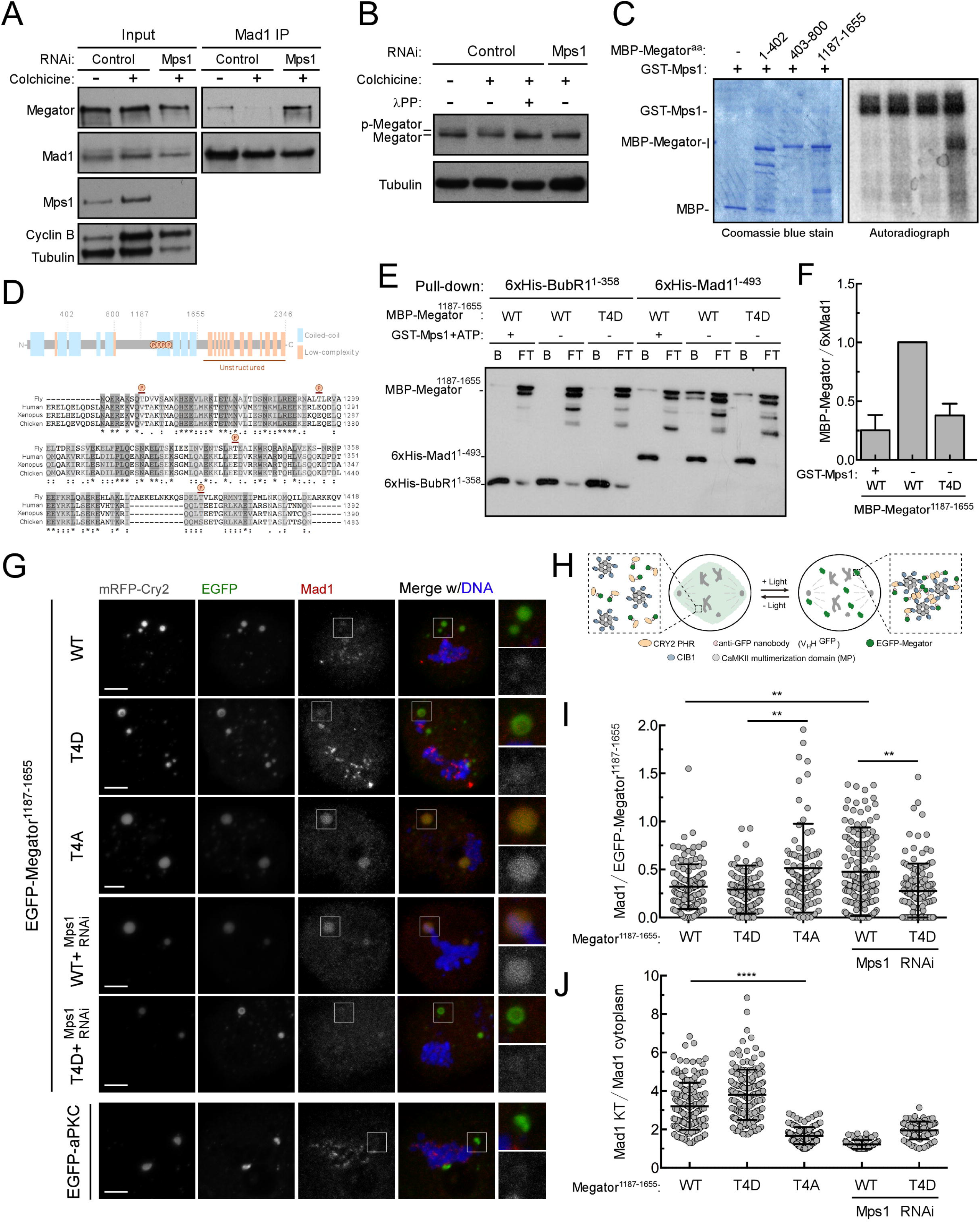
Msp1-mediated phosphorylation of Megator disrupts its interaction with Mad1. (A) Immunoprecipitates (IP) of Mad1 from lysates of asynchronous (-colchicine) and mitotically-enriched (+colchicine) S2 cultures incubated in the presence or absence of Mps1 RNAi for 120 hours. Colchicine (30μM) was incubated for 10 hours and MG132 (20μM) was added 4 hours prior to cell lysis. Mad1 IPs and corresponding inputs were blotted for the indicated proteins. (B) Western blot analysis of Megator hyperphosphorylation in cell lysates from the same experimental conditions as in (A). To validate the slower migrating band as an hyperphosphorylated form of Megator, the lysate of control colchicine-incubated cells was treated with λ-phosphatase (λPP) for 1 hour. (C) *In vitro* kinase assay with the indicated recombinant fragments of MBP-Megator and GST-Mps1 in the presence of [γ-^32^P]ATP for 30 min. Phosphorylation was detected by autoradiography and protein levels visualized by coomassie blue staining. (D) Schematic representation of *Drosophila* Megator obtained from the Eukaryotic Linear Motif (ELM) resource and Clustal Omega (EMBL-EBI) local sequence alignment for the indicated Megator/Tpr orthologues. Amino acids in dark gray background are conserved and amino acids in light gray background have similar chemical properties. Symbols: * fully conserved residue;: conservation between groups of strongly similar properties;. conservation between groups of weakly similar properties. Residues phosphorylated by Mps1 were identified by mass spectrometry analysis after *in vitro* kinase assay. Phospho-sites (P) were identified with MaxQuant/Andromeda with a decoy FDR of 0.01 on peptide and site level. (E) Pull-downs of recombinant purified MBP-Megator^1187-1655/WT^, MBP-Megator^1187-1655/WT^ phosphorylated by GST-Mps1, or MBP-Megator^1187-1655/T4D^ by bead-immobilized 6xHis-Mad1^1-493^ or 6xHis-BubR1 ^1-358^ (negative control). Both beads (B) and flow-through (FT) were blotted for the indicated proteins. (F) Quantification of MBP-Megator binding to 6xHis-Mad1^1-493^ from pull-downs in (E). The graph represents the ratio between the chemiluminescence signal intensities of MBP-Megator and 6xHis-Mad1^1-493^ from two independent experiments. The values obtained for MBP-Megator^1187-1655/WT^ were set to 1. (G,H) Representative immunofluorescence images (G) and schematic representation (H) of EGFP-Megator^1187-1655^ clustering by light-activated reversible inhibition by assembled trap (LARIAT) in mitotic S2 cells. Fusion of CIB1 with the multimerization domain from CaMKIIα (MP) forms dodecamers in the cytoplasm. The CRY2 photolyase homology region (PHR) is fused with an anti-GFP nanobody that binds specifically to EGFP-Megator. Blue light triggers CRY2 oligomerization and binding to CIB1, thus trapping EGFP-Megator into multimeric protein clusters. In the dark, CRY2 reverts spontaneously to its ground state and the clusters disassemble. LARIAT-induced clusters of EGFP-aPKC were used as negative control. The insets display magnifications of the outlined regions. S2 cells were treated with colchicine (30μM) for 10 hours and MG132 (20μM) for 4 hours followed by a 30 minute period of blue light irradiation. Expression of LARIAT-modules, EGFP-Megator^1187-1655^ transgenes and EGFP-aPKC was induced for 24 hours prior immunofluorescence analysis. (I,J) Quantification of Mad1 levels at EGFP-Megator^1187-1655^ clusters (I) and at kinetochores (J). Mad1 fluorescence intensities at clusters were determined relative to GFP-Megator^1187-1655^ signal (N>114 clusters for each condition) and at kinetochores relative to Mad1 cytosolic signal (N>64 kinetochores for each condition). Data information: in (F), (I), and (J), data are presented as mean ± SD. Asterisks indicate that differences between mean ranks are statistically significant, ** p<0.01, **** p<0.0001, (Kruskal-Wallis, Dunn’s multiple comparison test). Scale bars: 5μm

### Recruitment of Mad1 to unattached kinetochores requires its dissociation from Megator

We then sought to examine the relevance of disengaging Mad1 from Megator in mitosis. With the LARIAT experiment we observed that Mad1 levels at EGFP-Megator^1178-1655^ clusters and its accumulation at unattached kinetochores are inversely correlated (Figure 2G-J). This suggests that retaining Mad1 associated with Megator during mitosis precludes its proper recruitment to kinetochores. To address this further, we generated S2 cell lines stably expressing full length versions of Megator phosphomutants tagged with EGFP and depleted the endogenous nucleoporin with RNAi targeting the transcript UTRs (Figure 3A and 4A,B). Following an induction period of 24 hours, all transgenes are expressed at endogenous levels and localize correctly at the NE of interphase cells (Figure 3B; Figure 4A,B). Expression of Megator^T4D^-EGFP fails to rescue Mad1 loss from NPCs caused by depletion of the endogenous protein, further confirming that phosphorylation of T1259, T1395, T1338 and T1390 inhibits Megator interaction with Mad1 (Figure 3A-C). Conversely, Megator^T4A^-EGFP is able to restore Mad1 association with interphase NPCs (Figure 3A-C) but impairs its proper recruitment to unattached kinetochores. Expression of EGFP-Megator^T4A^ in colchicine-treated cells results in a two-fold reduction of kinetochore-associated Mad1 levels when compared to Megator^WT-^EGFP cells (Figure 3D,E). An antibody that specifically recognizes the closed conformer of Mad2 (Fava et al., 2011) reveals a similar decrease in the amount of C-Mad2 at unattached kinetochores of EGFP-Megator^T4A^ cells (Figure 4C,D). As expected, knocking-down Mps1 abrogates Mad1 and C-Mad2 kinetochore localization in cells expressing Megator^WT-^EGFP (both reduced to ~20% relative to control Megator^WT-^ EGFP cells, Figure 3D,E and 4C,D). Strikingly, this is ameliorated by precluding Mad1 interaction with Megator. Cells expressing Megator^T4D^-EGFP are still partially competent in recruiting Mad1, and to some extent, C-Mad2 to unattached kinetochores upon depletion of Mps1 kinase (~70% of Mad1 levels and ~50% of C-Mad2 levels relative to control Megator^WT-^EGFP cells, Figure 3D,E and 4C,D). Collectively, these results strongly suggest that kinetochore recruitment of a significant fraction (~50%) of Mad1-C-Mad2 heterotetramers requires the dissociation of Mad1 from Megator driven by Mps 1-mediated phosphorylation of the latter.

**Figure 3:**
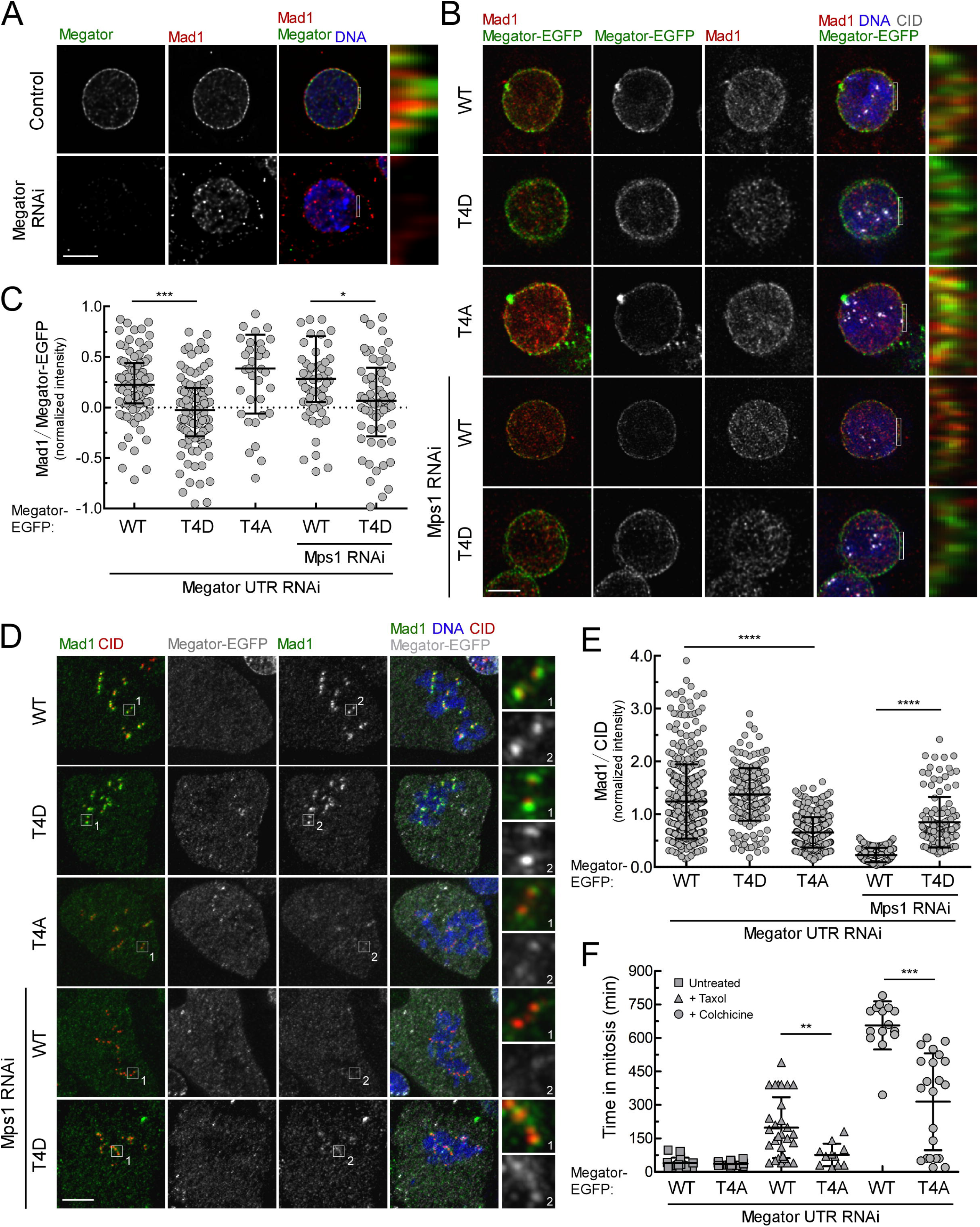
Recruitment of Mad1 to unattached kinetochores and robust SAC signaling require phosphorylation of Megator by Mps1. (A) Representative immunofluorescence images of Mad1 localization in control and Megator-depleted interphase cells. The insets display magnifications of the outlined regions. (B,C) Representative immunofluorescence images (B) and corresponding quantifications (C) of Mad1 at the nuclear envelope of interphase S2 cells depleted of endogenous Megator and expressing the indicated Megator-EGFP transgenes. When indicated, cultures were incubated in the presence of Mps1 RNAi for 120 hours. The insets display magnifications of the outlined regions. Mad1 fluorescence intensities were determined relative to Megator-EGFP signal (N ≥ 41 cells for each condition). (D,E) Representative immunofluorescence images (D) and corresponding quantification (E) of Mad1 at unattached kinetochores of S2 cells depleted of endogenous Megator and expressing the indicated Megator-EGFP transgenes. When indicated, cultures were incubated in the presence of Mps1 RNAi for 120 hours. To generate unattached kinetochores, cells were incubated with colchicine (30μM) and MG132 (20μM) for 30min prior to fixation. The insets display magnifications of the outlined regions. Mad1 fluorescence intensities were determined relative to CID signal (N ≥ 125 kinetochores for each condition). (F) Mitotic timings of S2 cells depleted of endogenous Megator and expressing the indicated Megator-EGFP transgenes under unperturbed conditions or upon addition of taxol (100nM) or colchicine (30μM) (N ≥ 11 cells for each condition). Expression of Megator-EGFP transgenes in (B-F) was induced for 24 hours prior processing for immunofluorescence analysis or live cell imaging. Data information: in (C) data is presented as median with interquartile range; in (E) and (F) data is presented as mean ± SD. Asterisks indicate that differences between mean ranks are statistically significant, * p<0.05, ** p< 0.01, *** p<0.001, **** p<0.0001 (Kruskal-Wallis, Dunn s multiple comparison test in (C) and (E) and Student’s t-test in (F)). Scale bars: 5μm.

### Dissociation of Mad1 from Megator in mitosis is required for functional SAC signalling

To examine the importance of Mps1-mediated phosphorylation of Megator for SAC signalling, we monitored by live-cell imaging the mitotic progression of Megator phosphomutants and assessed their capacity to arrest in mitosis when incubated with spindle poisons (Figure 3F). In asynchronous cultures, Megator^T4A^-EGFP cells progressed slightly faster from NEB to anaphase onset (~36 min) than cells expressing Megator^WT^-EGFP (~39 min), suggesting that the SAC might be partially compromised in the phosphodefective mutant. In line with SAC proficiency, cells expressing Megator^WT^-EGFP significantly delayed the transition to anaphase in the presence of colchicine (~640 min) or taxol (~180 min). In contrast, Megator^T4A^-EGFP cells, although able to exhibit some mitotic delay in response to unattached kinetochores (~360 min in colchicine) or decreased microtubules dynamics (~87 min in taxol), failed to maintain this to the same time extent as Megator^WT^-EGFP cells (Figure 3F). These results indicate that preventing the phosphorylation of Megator on T1259, T1395, T1338 and T1390 results in a weakened SAC function, which correlates with the observed reduction (~50%) in Mad1 and C-Mad2 levels at unattached kinetochores. Thus, we reason that phosphorylation of this patch of threonine residues by Mps1 kinase is required to release Mad1 from Megator to provide the kinetochore with sufficient amount of Mad1-C-Mad2 template that ensures a robust SAC response.

### Constitutive phosphorylation of Megator reduces C-Mad2 levels at kinetochores and compromises SAC strength

Mad1 and Mad2 form a highly stable complex *in vitro* (Sironi et al., 2002; De Antoni et al., 2005; Vink et al., 2006) and interact with each other throughout the cell cycle (Chen et al., 1998; Chung and Chen, 2002; Fava et al., 2011; Schweizer et al., 2013). Interestingly, although Megator^T4D^-EGFP cells can recruit Mad1 to unattached kinetochores, we observe an evident decline in the accumulation of C-Mad2 compared to cells expressing Megator^WT^-EGFP (Figure 4C,D). This phenotype is highly reminiscent of that observed in cells depleted of Megator (Figure 4F,G and S2D-F), which in mammals has been attributed to increased proteolytic degradation of Mad1 and Mad2 (Schweizer et al., 2013). Although we confirm in *Drosophila* cells a significant reduction in the total levels of Mad1 and Mad2 following Megator depletion (Figure 4A,B and S2D), these are partially rescued to a similar extent regardless the Megator-EGFP transgene expressed (Figure 4A,B). This argues against altered Mad1 and Mad2 proteostasis as the main underlying cause for deficient C-Mad2 accumulation at kinetochore of Megator^T4D^-EGFP cells. Instead, we envisage that Megator/Tpr might function *in vivo* as a scaffold for the assembly of the Mad1-C-Mad2 complex before its targeting to kinetochores. Kinetochore loading of C-Mad2 is therefore expected to occur inefficiently if Mad1 fails to bind Megator, as in interphase cells where the endogenous nucleoporin is depleted or replaced by the phosphomimetic version of the protein. In line with limited C-Mad2 at kinetochores, cells expressing Megator^T4D^-EGFP failed to arrest in mitosis in response to colchicine as efficiently as Megator^WT^-EGFP cells (Figure 4E). This is indicative of a weakened SAC function, which is also observed in parental S2 cells and human cultured cells respectively depleted of Megator (Figure 4H and S2G) or Tpr (Schweizer et al., 2013; Rodriguez-Bravo et al., 2014). Hence, while Mad1 dissociation from Megator triggered during mitotic entry efficiently endorses Mad1-C-Mad2 to unattached kinetochores, abrogating Mad1-Megator interaction during interphase is detrimental for SAC signalling as it possibly compromises the assembly of Mad1-C-Mad2 heterotetramers (Schweizer et al., 2013) and the formation of pre-mitotic MCC (Rodriguez-Bravo et al., 2014).

**Figure 4:**
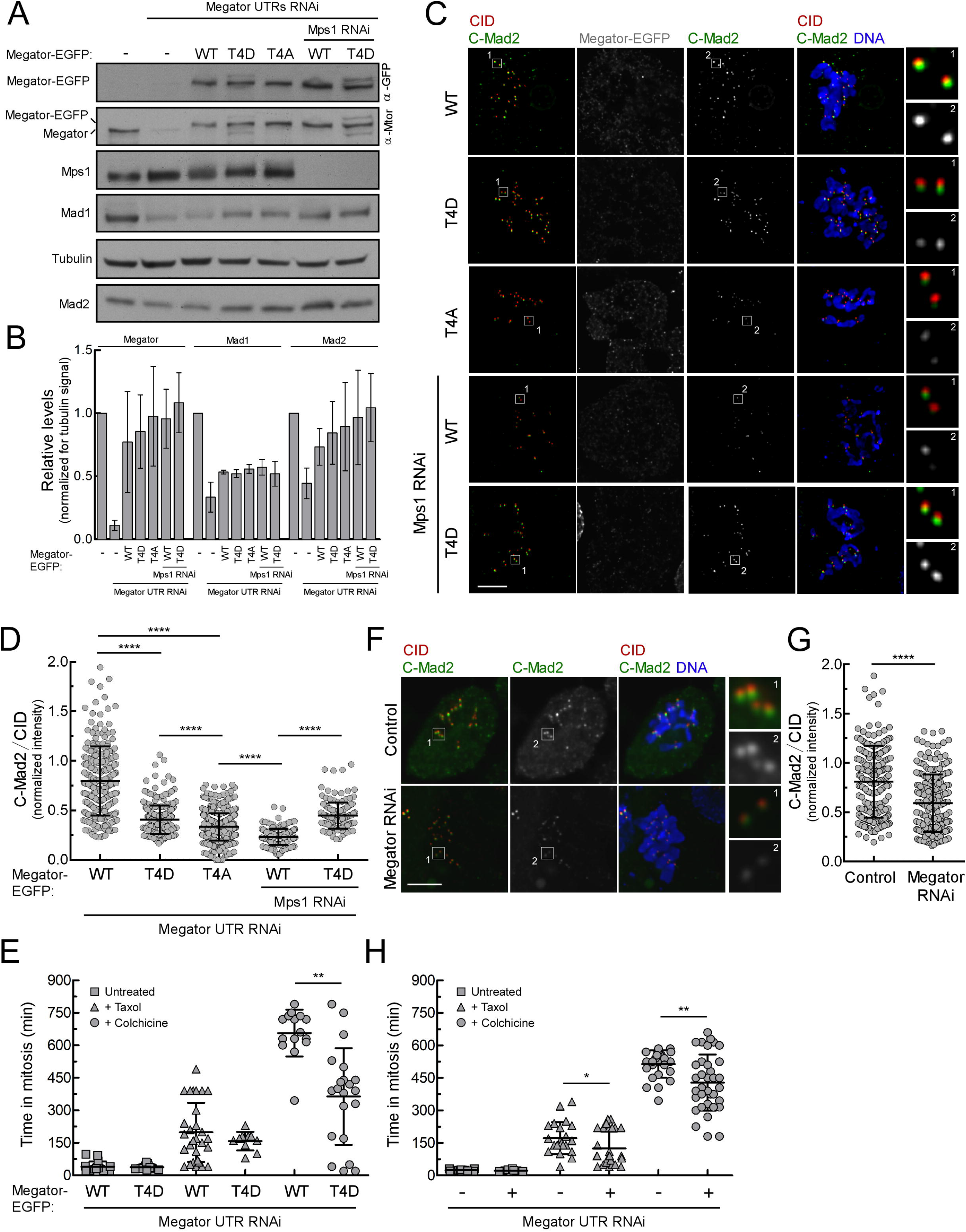
Constitutively impaired Mad1-Megator interaction reduces C-Mad2 levels at kinetochores and the strength of SAC signalling. (A,B) Representative western blots (A) and corresponding quantifications (B) of Megator, Mad1 and Mad2 protein levels in lysates from control or Megator-depleted S2 cells expressing the indicated Megator-EGFP transgenes. When indicated, cultures were incubated in the presence of Mps1 RNAi for 120h. The chemiluminescence signal intensities of Megator, Mad1 and Mad2 were determined relative to tubulin signal. The graph represents the quantification of relative protein levels from at least two independent experiments. The mean value obtained for control parental cells was set to 1. (C,D) Representative immunofluorescence images (C) and corresponding quantifications (D) of C-Mad2 at unattached kinetochores of S2 cells depleted of endogenous Megator and expressing the indicated Megator-EGFP transgenes. When indicated, cultures were incubated in the presence of Mps1 RNAi for 120 hours. To generate unattached kinetochores, cells were incubated with colchicine (30μM) and MG132 (20μM) for 30min prior to immunofluorescence analysis. The insets display magnifications of the outlined regions. C-Mad2 fluorescence intensities were determined relative to CID signal (N ≥ 148 kinetochores for each condition). (E) Mitotic timings of S2 cells depleted of endogenous Megator and expressing the indicated Megator-EGFP transgenes under unperturbed conditions or upon addition of taxol (100nM) or colchicine (30μM) (N ≥ 10 cells for each condition). Mitotic timings of S2 cells expressing Megator^WT^-EGFP are the same as used in Figure 3F. (F,G) Representative immunofluorescence images (F) and corresponding quantifications (G) of C-Mad2 levels at unattached kinetochores of control or Megator-depleted S2 cells treated with colchicine (30μM) and MG132 (20μM) for 30min. The insets display magnifications of the outlined regions. C-Mad2 fluorescence intensities were determined relative to CID signal (N ≥ 224 kinetochores for each condition). (H) Mitotic timings of control and Megator-depleted S2 cells under unperturbed conditions or upon addition of taxol (100nM) or colchicine (30μM) (N ≥ 8 cells for each condition). Expression of Megator-EGFP transgenes in (A-E) was induced for 24 hours prior processing for immunofluorescence analysis or live cell imaging. Data information: in (B) data is presented as mean ± SEM; in (D), (E), (G), and (H), data are presented as mean ± SD. Asterisks indicate that differences between mean ranks are statistically significant ****p<0.0001, (Kruskal-Wallis, Dunn s multiple comparison test in (D) and Student’s t-test in (E), (G) and (H). Scale bars: 5μm.

### Precluding Mad1 from binding to Megator rescues chromosome mis-segregation and aneuploidy in neuroblasts and intestinal stem cells depleted of Mps1

The results so far demonstrate that Mps1 controls Mad1 kinetochore recruitment in part by abolishing its interaction with Megator. We next assessed whether this mechanism occurs *in vivo* and its relevance for genomic stability. We prevented Mps1 kinetochore localization in *Drosophila* 3^rd^ instar larval neuroblasts and examined their capacity to recruit Mad1 to unattached kinetochores. For that, we resorted to *mps1*-null mutant flies (*ald^G4422^*) expressing a truncated version of Mps1 that lacks the N-terminus domain (gEGFP-Mps1-C^term^) required for kinetochore targeting (Althoff et al., 2012; Conde et al., 2013). As expected, Mad1 inability to localize at kinetochores of colchicine-treated *ald^G442^* neuroblasts was largely rescued by the expression of gEGFP-Mps1-WT (Figure S2A-C). Importantly, neuroblasts expressing the gEGFP-Mps1-C^term^ transgene under control of *mps1* promoter were still able to partially recruit Mad1 to unattached kinetochores, accumulating up to 50% of the levels detected in control *w^1118^* flies (Figure S2A-C). This confirms *in vivo* a kinetochore-extrinsic role for Mps1 in Mad1 kinetochore localization.

Collectively, our results indicate that non-kinetochore Mps1 abolishes the interaction between Megator and Mad1 in mitosis, which would otherwise preclude efficient recruitment of Mad1-C-Mad2 to unattached kinetochores. This rationale is further supported *in vivo* by the observation that RNAi-mediated repression of Megator extensively restores Mad1 kinetochore recruitment in a *ald^G4422^* genetic background (Figure 5A,B). Importantly, this concomitantly rescues the aneuploidy that is caused by loss of Mps1 activity (Figure 5A). Depletion of Megator from *ald^G4422^* neuroblasts led to a striking decrease in the frequency of mitotic figures exhibiting aneuploid karyotypes (~70% in *ald^G4422^ vs* ~30% in *ald^G4422^+Megator* RNAi; Figure 5A). Given the evident recovery in genomic stability, we hypothesized that depletion of Megator improves the fidelity of chromosome segregation in cells devoid of Mps1 activity. To test this, we monitored by live imaging the mitotic progression of unperturbed (no drugs) larval neuroblasts (Figure 5C-E). Consistent with Mps1 and Megator roles in SAC signalling, neuroblasts depleted of either protein progressed faster through mitosis (~5 min), when compared to *w^1118^* controls (~7.3 min). Interestingly, although *UAS-MegatorRNAi* and *ald^G4422^* neuroblasts reveal indistinguishable mitotic timings, the frequency of anaphases with lagging chromosomes is dramatically higher in *mps1-null* mutants (Figure 5C-E). Importantly, depletion of Megator significantly restores the accuracy of segregation in *ald^G4422^* neuroblasts, albeit failing to extend the time from NEB to anaphase onset (Figure 5C-E). A similar defect in SAC function was observed in cultured S2 cells co-depleted of Mps1 and Megator. Although proficient in Mad1 kinetochore recruitment (Figure S2E,F), these cells failed to arrest in mitosis when challenged with colchicine (Figure S2G), as would be expected from compromised Mps1-dependent phosphorylation of Mad1, and consequently, from limited C-Mad2-Cdc20 interaction (Faesen et al., 2017; Ji et al., 2017). Hence, in *Drosophila* neuroblasts undergoing unperturbed mitosis, kinetochore-associated Mad1 is able to efficiently safeguard anaphase fidelity and chromosomal euploidy independently of its SAC function. This is in line with previous studies reporting a role for Mad1 in preventing merotely through pathways that are uncoupled from its interaction with Mad2 and SAC signalling (Emre et al., 2011; Akera et al., 2015). We then tested whether a similar improvement in genomic stability also occurs in adult tissues. We have recently shown that inducing aneuploidy in intestinal stem cells (ISCs) through depletion of Mps1 results in severe intestinal dysplasia (Resende et al., 2018, Figure S3A). Here, we confirmed an increased proliferation of ISCs and enteroblasts (EBs) following *EsgGAL4-driven* expression of *UAS-Mps1RNAi* (Figure S3B,C). Importantly, we found that co-expression of *UAS-MegatorRNAi* supresses this dysplastic phenotype, thus suggesting a rescue in the levels of aneuploidy in ISCs as observed for larval neuroblasts. Collectively, these results strongly support that Mad1 dissociation from its nuclear pore receptor represents a critical event for efficient kinetochore localization, for the fidelity of chromosome segregation and consequently, for genome stability *in vivo.*

**Figure 5.**
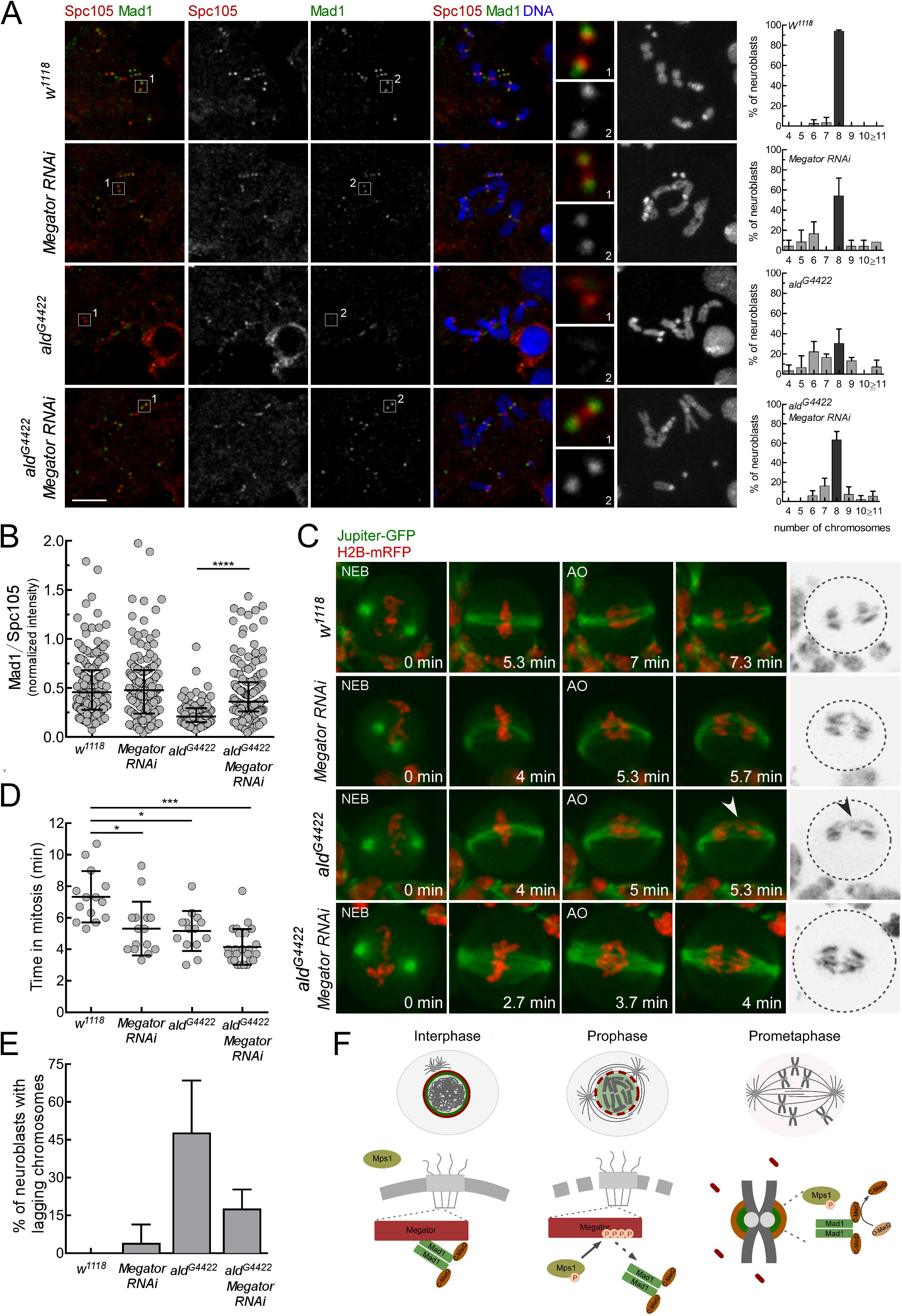
Depletion of Megator restores Mad1 kinetochore recruitment and mitotic fidelity in *Drosophila mps1-null* neuroblasts. (A,B) Representative immunofluorescence images with ploidy histograms (A) and corresponding quantifications (B) of Mad1 levels at unattached kinetochores of *w^1118^, InscGal4>UAS-MegatorRNAi, ald^G4422^* or *ald^G4422^;InscGal4>UAS-MegatorRNAi* neuroblasts treated with colchicine (50μM) for 1.5 hours. The insets display magnifications of the outlined regions. Mad1 fluorescence intensities were determined relative to Spc105 signal. (N>91 kinetochores for each condition) (C-E) Mitotic progression (C), mitotic timing (D) and percentage of anaphases with lagging chromosomes (E) of *w^1118^, InscGal4>UAS-MegatorRNAi, ald^G4422^* or *alcr^4422^;InscGal4> UAS-MegatorRNAi* neuroblasts co-expressing Jupiter-GFP and H2B-mRFP. Mitotic progression was monitored through time-lapse microscopy and the mitotic timing was defined as the time cells spent from nuclear envelope breakdown (NEB) to anaphase onset (AO) (N 14 neuroblasts for each condition from at least two independent experiments). The arrowhead in (C) points to a lagging chromosome. (F) Proposed model for the control of Mad1 subcellular redistribution during the G2/M transition. In interphase, inactive Mps1 (unphosphorylated T-loop) is retained in the cytoplasm and Mad1-C-Mad2 complexes are docked at the nucleoplasmic side of NPCs through Mad1 binding to Megator. During prophase, active Mps1 (phosphorylated T-loop) becomes detectable in the nucleus and is now able to phosphorylate Megator. This disrupts the nucleoporin interaction with Mad1, hence ensuring timely release of Mad1-C-Mad2 from NPCs. Dissociation from Megator enables Mad1-C-Mad2 to efficiently accumulate at prometaphase kinetochores and instate robust SAC signalling. Data information: in (A), (B), (C), and (E), data are presented as mean ± SD. Asterisks indicate that differences between mean ranks are statistically significant, *p < 0.05, *** p<0.001, ****p<0.0001 (Kruskal-Wallis, Dunn’s multiple comparison test). Scale bars: 5μm.

Our biochemical, cellular and *in vivo* data concur to demonstrate that Mps1 activity at NPCs early in prophase sets the stage to enable appropriate recruitment of Mad1 by unattached/prometaphase kinetochores (Figure 5F). Phosphorylation of Megator by Mps1 abrogates the nucleoporin interaction with Mad1, which we find to be essential for kinetochore localization of Mad1-C-Mad2 to levels required to sustain robust SAC signalling and accurate chromosome segregation. We also show that dissociation of Mad1 from NPCs is prevented during interphase by nuclear exclusion of Mps1 and decreased phosphorylation of its activating T-loop. This is most likely important to facilitate Mad1-C-Mad2 interaction and the assembly of MCC prior to kinetochore maturation and SAC activation (Rodriguez-Bravo et al., 2014; Kim et al., 2018). Together, these observations establish that a key function of non-kinetochore Mps1 is to coordinate Mad1 subcellular localization with cell cycle progression, so that both nuclear pores in interphase and kinetochores in mitosis generate anaphase inhibitors that preserve genomic stability (Figure S4).

## MATERIALS AND METHODS

### S2 cell cultures, RNAi-mediated depletion and drug treatments

The *Drosophila* S2-DGRC cell line (#stock6) was acquired from the Drosophila Genomics Resource Center, Indiana University and was not independently authenticated. The cell lines were routinely tested negative for mycoplasma contamination. Cell cultures, RNAi synthesis and RNAi treatment were performed as previously described (Conde et al., 2013). At selected time points, cells were collected and processed for immunofluorescence, time-lapse microscopy, immunoblotting or immunoprecipitation. When required, cells were subjected to several drug treatments before being collected and processed for the desired analysis. To promote microtubule depolymerisation, cells were incubated with 30 μM colchicine (Sigma–Aldrich, St. Louis, MO) for 30 minutes – 24 hours. To decrease microtubule dynamics cells were incubated with 100 nM taxol (Sigma-Aldrich). When required, 20μM MG132 (Calbiochem, San Diego, CA) were added to inhibit the proteasome. For experiments in Figure 1D,E and Figure S1C, cells were incubated with 10μM of Leptomycine B (Sigma-Aldrich) for 3 hours to block Crm1-mediated nuclear export.

### Constructs and S2 cells transfection

Recombinant plasmids pHWG[blast]-Megator^WT^, pHWG[blast]-Megator^T4A^, pHWG[blast]-Megator^T4D^, pHGW[blast] -Megator^1187-1655/WT^, pHGW[blast] -Megator^1187-1655/T4A^, pHGW[blast]-Megator^1187-1655/T4D^ and pHGW[blast]-Mps1^WT^ were generated using the Gateway Cloning System (Invitrogen). Megator, Megator^1187-1655^ or Mps1 cDNAs were amplified by PCR and inserted into modified versions of pENTR-entry vector through FastCloning (Li et al., 2011). To generate pENTR-Megator^1187-1655/T4A^ and pENTR-Megator^1187-1655/T4D^ codons corresponding to T1259, T1302, T1338 and T1390 of pENTR-Megator^1187-1655^ were converted either to codons for alanine (A) or aspartate (D) respectively, by several cycles of site-directed mutagenesis with primers harbouring the desired mutations. To generate pENTR-Megator^T4A^ and pENTR-Megator^T4D^, the fragment corresponding to amino acids 1187-1655 on pENTR-Megator^WT^ was replaced by Megator^1187-1655/T4A^ or Megator^1187-1655/T4D^ PCR products, respectively through FastCloning (Li et al., 2011). PCR reactions were performed using Phusion polymerase (New England Biolabs). PCR products were digested with DpnI restriction enzyme (New England Biolabs), used to transform competent bacteria and selected for positives. Subsequently, pENTR-Megator^1187-1655^ constructs and pENTR-Mps1 were recombined with pHGW[blast] (blasticidin^R^; N-terminal EGFP tag), and pENTR-Megator constructs with pHWG (blasticidin^R^; C-terminal EGFP tag) using Gateway LR Clonase II (Invitrogen), according to the manufacturer’s instructions. pHGW-Mps1^WT^-NLS was produced by PCR amplification of pHGW-Mps1^WT^ with primers harbouring SV40 large T-antigen nuclear localization signal sequence. pHGW-Mps1^KD^-NLS was produced by site-directed mutagenesis of pHGW-Mps1^WT^-NLS with primers harbouring the mutation to convert D478 to A478. PCR reactions were performed with Phusion polymerase, followed by digestion with DpnI restriction enzyme (New England Biolabs). The constructs H2B-mCherry, H2B-GFP, mCherry-α-Tubulin, Mad1-EGFP, pHW-CIB-MP-HRW-CRY2-V_H_H, and pHGW-aPKC have been previously described (Conde et al., 2013; Moura et al., 2017; Osswald et al., 2019). Plasmids were transfected into S2 cells using Effectene Transfection Reagent (Qiagen), according to the manufacturer’s instructions. Transiently expressing cells were harvested 4-5 days after transfections. Stable cell lines were obtained by selection in medium with 25 μg/mL blasticidin. To induce expression of pHW-CIB-MP-HRW-CRY2-V_H_H, pHGW[blast] or pHWG[blast] constructs cells were incubated for 30min at 37°C 24 hours prior to processing. Cells transfected with pHW-CIB-MP-HRW-CRY2-V_H_H were maintained in the dark until processing.

### Live cell imaging

Live analysis of mitosis was performed in S2 cell lines and neuroblasts expressing the indicated constructs. S2 cells were plated on glass bottom dishes (MatTek) coated with Concanavalin A (0.25 mg/mL; Sigma-Aldrich). Third-instar larvae brains were dissected in PBS and mounted in PBS between coverslips of different sizes. The preparation was squashed and sealed with Halocarbon oil 700 (Sigma-Aldrich). 4D datasets were collected at 25°C with a spinning disc confocal system (Revolution; Andor) equipped with an electron multiplying charge-coupled device camera (iXonEM+; Andor) and a CSU-22 unit (Yokogawa) based on an inverted microscope (IX81; Olympus). Two laser lines (488 and 561 nm) were used for near-simultaneous excitation of EGFP and mCherry or RFP. The system was driven by iQ software (Andor). Time-lapse imaging of z stacks with 0.8 μm steps for S2 cells and 0.5μm for neuroblasts were collected and image sequence analysis, video assembly and fluorescence intensities quantification performed using ImageJ software. Quantification of Mad1-EGFP and Megator-EGFP levels at the nuclear envelope and mCherry tubulin at the nucleus was performed on single Z stacks from images acquired with fixed exposure settings. Mad1-EGFP and Megator-EGFP intensities at the nuclear envelope were determined for each time point (t), using the following formula:

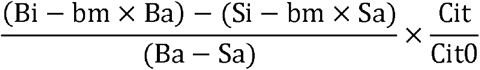

Bi-integrated density of a ROI harbouring the nucleus (including outer nuclear membrane); Ba – area of the ROI harbouring the nucleus; Si – integrated density of a ROI encompassing the nucleoplasm; Sa – area of the ROI harbouring the nucleoplasm; bm-mean intensity a ROI outside the cell (background); Cit – integrated density of a ROI harbouring the cell at time t; Cit0-integrated density of a ROI harbouring the cell on the first frame. mCherry-Tubulin intensities in the nucleus were determined for each time point (t), using the following formula:

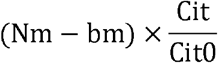

Nm- mean intensity of a ROI inside the nucleus, bm-mean intensity of the background, Cit-integrated density of a ROI harbouring the cell at time t; Cit0-integrated density of a ROI harbouring the cell on the first frame. The changes in fluorescence intensity over time were plotted as normalized signal relative to the mean signal measured before NEB.

### Immunofluorescence analysis

For immunofluorescence analysis of S2 cells, 10^5^ cells were centrifuged onto slides for 5 min, at 1500 rpm (Cytospin 2, Shandon). For LARIAT experiments, cells were irradiated with blue light for 30 min prior to centrifugation. Afterwards, cells were fixed in 4% paraformaldehyde in PBS for 12min and further extracted for 8min with 0.1% Triton X-100 in PBS. Alternatively, cells were simultaneously fixed and extracted in 3.7% formadehyde (Sigma), 0.5 % Triton X-100 in PBS for 10min followed by three washing steps of 5 min with PBS-T (PBS with 0,05% Tween20). For immunofluorescence analysis of *Drosophila* neuroblasts, third-instar larval brains were dissected in PBS and incubated with 50μM colchicine for 1.5h. The brains were after fixed in 1.8% formaldehyde (Sigma-Aldrich) and 45% glacial acetic acid for 5min, squashed between slide and coverslip and immersed in liquid nitrogen. Subsequently, coverslips were removed, the slides were incubated in cold ethanol for 10 min and washed in PBS with 0.1% Triton X-100. Immunostaining was performed as previously described (Moura et al., 2017). Fixation and immunostaining of intestines from 20 day adult flies were performed as previously described (Resende et al., 2018). Images were collected in a Zeiss Axio Imager microscope (Carl Zeiss, Germany) or in a Leica TCS II scanning confocal microscope (Leica Microsystems). For immunofluorescence quantification, the mean pixel intensity was obtained from raw images acquired with fixed exposure acquisition settings. Fluorescence intensities at the nuclear envelope were obtained from single Z stack projections. The nuclear envelope was defined based on Megator or Megator-EGFP staining, by subtracting a ROI containing the nucleoplasm to a ROI harboring the entire nucleus (outer membrane) after subtraction of background intensities estimated from regions outside the cell. Mad1 fluorescence intensities were determined relative to Megator or Megator-EGFP. Fluorescence intensities of LARIAT-mediated clustered proteins and kinetochore proteins were obtained from maximum projected images. For Mad1 and EGFP-Megator^1187-1655^, the fluorescence intensities were quantified for individual clusters selected manually by mRFP-Cry2 staining. After subtraction of background intensities estimated from regions inside of the cell with no clusters, the intensity of Mad1 was determined relative to Megator-EGFP signal. For kinetochore proteins the fluorescence intensity was quantified for individual kinetochores selected manually by Mad1, CID or Spc105 staining. The size of the ROI was predefined so that each single kinetochore could fit into. After subtraction of background intensities, estimated from regions outside the cell, the intensity of the proteins was determined relative to cytoplasmic Mad1, CID or Spc105.

### S2 cell lysates, immunoprecipitation, and western blotting

For *Drosophila* brain lysates, at least 10 third-instar larvae brains were dissected in PBS, transferred to Laemmli Buffer (4% SDS, 10% β-mercaptoetanol, 0.125M Tris-HCl, 20% glycerol and 0.004% bromophenol blue) and boiled at 95°C for 5min. S2 cell lysates for immunoprecipitation and western blot analysis were obtained from non-transfected S2 cells or S2 cells expressing Megator-EGFP transgenes treated with colchicine and MG132 when indicated. For western blot of total S2 cell lysates, 10^6^ cells were harvested through centrifugation at 5000rpm for 10min. The resulting pellet was resuspended in Laemmli sample buffer and boiled at 95°C for 5min.

For immunoprecipitation assays, cells were harvested through centrifugation at 5000 rpm for 10 min at 4°C and afterwards washed with 2mL PBS supplemented with 1x protease inhibitors cocktail (Roche, Basel, Switzerland). Cell pellet was resuspended in lysis buffer (150 mM KCl, 75 mM HEPES, pH 7.5, 1.5 mM EGTA, 1.5 mM MgCl2, 15% glycerol, 0.1% NP-40, 1x protease inhibitors cocktail (Roche) and 1x phosphatase inhibitors cocktail 3 (Sigma)) before freezing disruption in liquid nitrogen. Cell lysates were then clarified through centrifugation at 10000 rpm for 10 min at 4°C and concentration determined measuring Absorbance at 280nm in the Nanodrop 1000 (ThermoFisher). Lysates containing 800μg of protein in a total of 400μl lysis buffer were pre-cleared by incubation with 15μl of Protein A magnetic beads (New England Biolabs) for 1h at 4°C under agitation. Pre-cleared extracts were incubated with rabbit anti-Mad1 (1:100) overnight, at 4°C under agitation. Afterwards, the mixture was incubated with 40μl of Protein A magnetic beads for 1h30 min, at 4°C with agitation. Magnetic beads were collected and washed 4 times with 500μL of lysis buffer. Magnetic beads were resuspended Laemmli sample buffer and boiled at 95°C for 5 min. To confirm protein hyperphosphorylation status, 50μg of mitotic cell lysates were treated with 400U of λ-phosphatase (New England Biolabs) at 30°C for 1 hour in a total volume of 50μl PMP phosphatase buffer (50 mM HEPES pH 7.5, 100 mM NaCl, 2 mM DTT, 0.01% Brij 35, 1 mM MnCl_2_; New England Biolabs).

Samples were resolved by SDS-PAGE and transferred to a nitrocellulose membrane, using the iBlot Dry Blotting System (Invitrogen) according to the manufacturer’s instructions. Transferred proteins were confirmed by Ponceau staining (0.25% Ponceau S in 40% methanol and 15% acetic acid). The membrane was blocked for 1 hour at room temperature with 5% powder milk prepared in PBST and subsequently incubated with primary antibodies diluted in blocking solution overnight at 4°C under agitation. Membranes were washed three times for 10min with PSBT and incubated with secondary antibodies (diluted in blocking solution) for 1 hour at room temperature with agitation. Secondary antibodies conjugated to Horseradish peroxidase (Santa Cruz Biotechnology) or VeriBlot for IP Detection Reagent (HRP) (Abcam, ab131366) were used according to the manufacturer’s instructions. Blots were developed with ECL Chemiluminescent Detection System (Amersham) according to manufacturer’s protocol and detected on X-ray film (Fuji Medical X-Ray Film). When required proteins were resolved in 4-20% Mini-PROTEAN^®^ TGX Precast Gel (BioRad) and transferred to nitrocellulose membrane overnight in 48mM Tris, 39mM glycine, 0.037% SDS, 20% metanol, pH=8.3, at 20V, 4°C.

### Production and purification of recombinant proteins

To generate 6xHis-Mad 1^1-493^ and 6xHis-BubR1 ^1-358^ constructs for expression in bacteria, PCR products with the coding sequence were cloned into NdeI/XhoI or SalI/XhoI sites of pET30a (+) vector (Novagen, Darmstadt, Germany), respectively. TOP10 competent cells were transformed and selected for positives. The recombinant construct was used to transform BL21-star competent cells and protein expression induced with 0.05mM IPTG at 15°C, overnight. Cells were harvested and lysed in bacterial lysis buffer (50mM NaH_2_PO_4_, 300mM NaCl, 10mM imidazole, pH=8.0) supplemented with 1mM PMSF (Sigma), 0.4mg/ml Lysozyme (Sigma), sonicated and clarified by centrifugation at 4°C. Recombinant 6xHis-Mad1^1-493^, 6xHis-BubR1 ^1-358^ were purified with Novex Dynabeads (Invitrogen) in bacterial lysis buffer.

To generate recombinant MBP-Megator fragments (a.a. 1-402; 403-800; 1187-1655), PCR products harboring the coding sequences for fragments 1-402, 403-800 and 1187-1655 of

Megator were cloned into pMal-c_2_ (New England Biolabs) vector. Megator ^1187-1655/T4A^ and Megator ^1187-1655/T4D^ were inserted into pMal-c2 vector through FastCloning (Li et al., 2011) using Phusion Polymerase (New England Biolabs). These constructs were used to transform TOP10 competent bacteria and cells were selected for the incorporation of plasmids. The selected recombinant constructs were used to transform BL21-star competent cells and protein expression induced with 0.05 mM IPTG at 15°C, overnight. Pellets of these cultures were lysed in column buffer (200 mM NaCl, 20 mM Tris-HCl, 1 mM EDTA, 1mM DTT, pH=7.4) supplemented with 1% Triton X100 (Sigma), 1mM PMSF (Sigma), and 0.4 mg/ml of lysozyme (Sigma), sonicated and clarified by centrifugation at 4°C. Recombinant MBP-Megator fragments were purified with amylose magnetic beads (New England Biolabs) and eluted in Column Buffer supplemented with 10mM Maltose. The purified recombinant proteins (eluted or bound to magnetic beads) were resolved by SDS-PAGE and their relative amounts determined after Coomassie blue staining. Similar amounts of protein were used in the subsequent assays.

### *In vitro* kinase assays, mass spectrometry analysis and pull-down assays

For *in vitro* kinase assays, recombinant fragments of MBP-Megator were incubated with 0.05μg HsMps1/TTK (SignalChem, Richmond, Canada) in a total volume of 30μl kinase reaction buffer (5 mM MOPS pH 7.2, 2.5 mM β-glycerol-phosphate, 5 mM MgCl2, 1 mM EGTA, 0.4 mM EDTA, 0.25 mM DTT, 100 μM ATP and supplemented with 1× phosphatase inhibitors cocktail 3 (Roche). Reactions were carried out at 30°C for 30 min, and analysed by autoradiography, subjected to mass spectrometry analysis or used in pull-down assays. For detection of ^32^P incorporation, the kinase reaction buffer was supplemented with 10 μ Ci[*γ* – ^32^P] ATP [3000Ci/mmol, 10mCi/mL] and the reaction was stopped by addition of Laemmli sample buffer, boiled for 5min at 95°C and resolved by SDS-PAGE. After drying at 80°C under vacuum, the gel was exposed to X-ray films (Fuji Medical X-Ray Film). For identification of phosphorylated residues, the reaction was stopped by addition of 6M Urea and subsequently analyzed by liquid chromatography coupled with mass spectrometry. Samples were digested with LysC/Trypsin and/or GluC and prepared for LC-MS/MS analysis as previously described (Rappsilber et al., 2007). Peptides (100ng) were separated on a Thermo ScientificTM EASY-nLC 1000 HPLC system (Thermo Fisher ScientificTM) for 1hour from 5-60% acetonitrile with 0.1% fromic acid and directly sprayed via a nanoelectrospray source in a quadrupole Orbitrap mass spectrometer (Q ExactiveTM, Thermo Fisher ScientificTM) (Michalski et al., 2011). The Q ExactiveTM was operated in data-dependent mode acquiring one survey scan and subsequently ten MS/MS scans (Olsen et al., 2007). Resulting raw files were processed with the MaxQuant software (version 1.5.2.18) using a reduced database containing only the proteins of interest and giving phosphorylation on serine, threonine and tyrosine as variable modification (Cox and Mann, 2008). A false discovery rate cut off of 1% was applied at the peptide and protein levels and the phosphorylation site decoy fraction.

For pull-down assays, 6xHis-Mad1^1-493^ or 6xHis-BubR1 ^1-358^ bound to Novex Dynabeads (Invitrogen) were incubated with the MPB-Megator^1187-1655^ constructs in a final volume of 50μL column buffer (250 mM NaCl, 20 mM Tris-HCl pH 7.4, 1 mM EDTA, 1 mM DTT, 0.05% Tween20 (Sigma) 1x protease inhibitors cocktail (Roche) and 1x phosphatase inhibitors cocktail 3 (Sigma-Aldrich) for 1h30min at room temperature with agitation. The magnetic beads (with bound protein) were collected and washed 3 times with 500μL column buffer, resuspended in Laemmli sample buffer and boiled at 95°C for 5 min. After removal of the magnetic beads, samples were resolved by SDS-PAGE and probed for proteins of interest through western blotting.

### Antibodies

The following primary antibodies were used for immunofluorescence studies: rat anti-CID (Rat4) used at 1:250, rabbit anti-phosphorylated Thr676-Mps1 (T676) (a gift from Geert Kops, (Jelluma et al., 2008) used at 1:2000, chicken anti-GFP (Abcam, ab 13970) used at 1:2000 for S2 cells and 1:1000 in neuroblasts, mouse anti-Megator (gift from Jørgen Johansen and Kristen Johansen, Qi et al., 2004, RRID:AB_2721935), used at 1:20, rabbit anti-Mad1 (Rb1, Conde et al., 2013) used at 1:2500 for S2 cells and 1:1000 for neuroblasts, mouse anti-C-Mad2 (Sigma), used at 1:50 for S2 cells and 1:25 for neuroblasts, rat anti-Spc105 used at 1:250, guinea pig anti-Mps1 (Gp15) (a gift from Scott Hawley, RRID:AB_2567774) used at 1:250, rabbit anti-phosphorylated ser10-Histone H3 (p-H3) (Milipore, Billerica, MA, RRID:AB_565299) used at 1:5000, rabbit anti-GFP (Molecular Probes) used at 1:5000 for *Drosophila* intestines. The following primary antibodies were used for western blotting studies: mouse anti-α-tubulin DM1A (Sigma-Aldrich, RRID:AB_477593) used at 1:10000; rabbit anti-Cyclin B (gift from C. Lehner) used at 1:10000, guinea pig anti-Mps1 (Gp15) (a gift from Scott Hawley, RRID:AB_2567774) used at 1:5000; mouse anti-Megator (gift from Jørgen Johansen and Kristen Johansen, Qi et al., 2004, RRID:AB_2721935) used at 1:100, rabbit anti-Mad1 (Rb1,(Conde et al., 2013) used at 1:2000, rabbit anti-Mad2 (Rb 1223) used at 1:100, mouse anti-MBP (New England Biolabs, RRID:AB_ 1559738), used at 1:5000; mouse anti His Tag (Milipore, 05-949) used at 1:2500.

### Fly stocks

All fly stocks were obtained from Bloomington Stock Center (Indiana, USA), unless stated otherwise. The mps1 mutant allele *ald^G4422^* has been described before (Conde et al., 2013). *Insc-GAL4* was used to drive the expression of *UAS-MegatorRNAi* and *UAS-Mad1RNAi* in neuroblasts from third-instar larvae brains. *EsgGAL4* was used to drive expression of *UAS-MegatorRNAi* and *UAS-Mps1RNAi* in ISCs and EBs from intestines of adult flies, as previously described (Resende et al., 2018). w^1118^ was used as wild-type control. Fly stocks harboring gEGFP-MPS1^WT^ and gEGFP-MPS1^325-630^ under control of Mps1 cis-regulatory region were kindly provided by Christian Lehner (Althoff et al., 2012).

### Statistical analysis

All statistical analysis was performed with GraphPad Prism V7.0f (Graph-Pad Software, Inc.).

## ACKNOWLEDGEMENTS

We thank Geert Kops (Hubrecht Institute, The Netherlands) for the phospho-specific Mps1 antibody, Thomas Maresca (University of Massachusetts Amherst, USA) for the Mad1-EGFP construct, Christian Lehner (University of Zurich, Switzerland) for the gEGFP-Mps1-WT and gEGFP-Mps1-C^term^ fly stocks, Eurico Morais-de-Sá (i3S/IBMC, Portugal) for sharing unpublished LARIAT constructs, Helder Maiato (i3S/IBMC, Portugal) for the Megator cDNA and Jørgen Johansen and Kristen Johansen (Iowa State University, USA) for the Megator antibody. We thank Ana Pinto, Nelson Leça and Margarida Moura (i3S/IBMC, Portugal) for technical help and critical reading of the manuscript.

This article is a result of the project Norte-01-0145-FEDER-000029 – Advancing Cancer Research: From basic knowledge to application, supported by Norte Portugal Regional Operational Programme (NORTE 2020), under the PORTUGAL 2020 Partnership Agreement, through the European Regional Development Fund (FEDER). MO is supported by a fellowship from the GABBA PhD program from the University of Porto, PD/BD/105746/2014. SC-S is supported by the FCT PhD fellowship SFRH/BD/136527/2018). JB is supported by an FCT PhD grant SFRH/BD/87871/2012. CC is supported by an FCT investigator position and funding (IF/01755/2014).

The authors declare no competing financial interests

## AUTHOR CONTRIBUTIONS

Mariana Osswald and Sofia Cunha-Silva performed most of the experiments with contributions from Jana Goemann, Luis M Santos and Carlos Conde. João Barbosa performed the live-imaging experiments with *Drosophila* neuroblasts. Pedro Resende performed the immunofluorescence analysis of *Drosophila* intestines. Tanja Bange performed the mass-spectrometry analysis of *in vitro* kinase assays. Mariana Osswald, Sofia Cunha-Silva, Claudio E Sunkel and Carlos Conde analysed the data. Mariana Osswald and Carlos Conde conceived the project. Carlos Conde designed the experiments, wrote the manuscript and coordinated the project.

---

**Figure S1:**
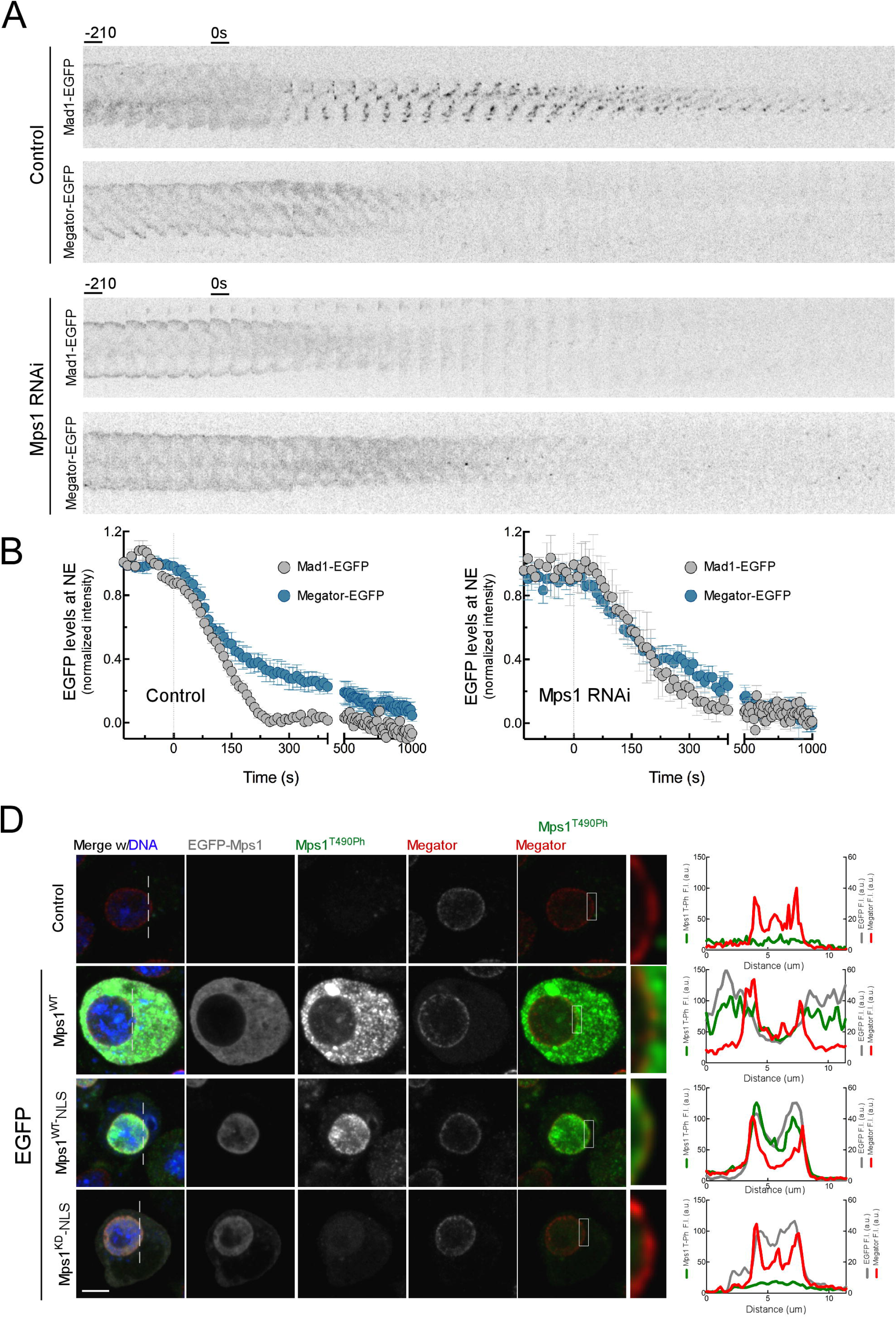
Additional information related to Figure 1. (A) Kymograph representations of Mad1-EFGP and Megator-EGFP localization pattern from movies in Figure 1A. (B) Data corresponding to the quantifications of nuclear envelope Mad1-EFGP and Megator-EGFP from Figure 1B plotted in the same graph for comparison purposes. (C) Representative immunofluorescence images of EGFP-Mps1, Mps1^T490Ph^ and Megator localization pattern in interphase control S2 cells and interphase S2 cells expressing EGFP-Mps1^WT^, EGFP-Mps1^WT^-NLS or EGFP-Mps1^KD^-NLS. The insets display magnifications of the outlined regions. Graphs represent the intensity profiles of GFP-Mps1, Mps1^T490Ph^ and Megator signal along the dotted lines. Scale bar: 5μm

**Figure S2.**
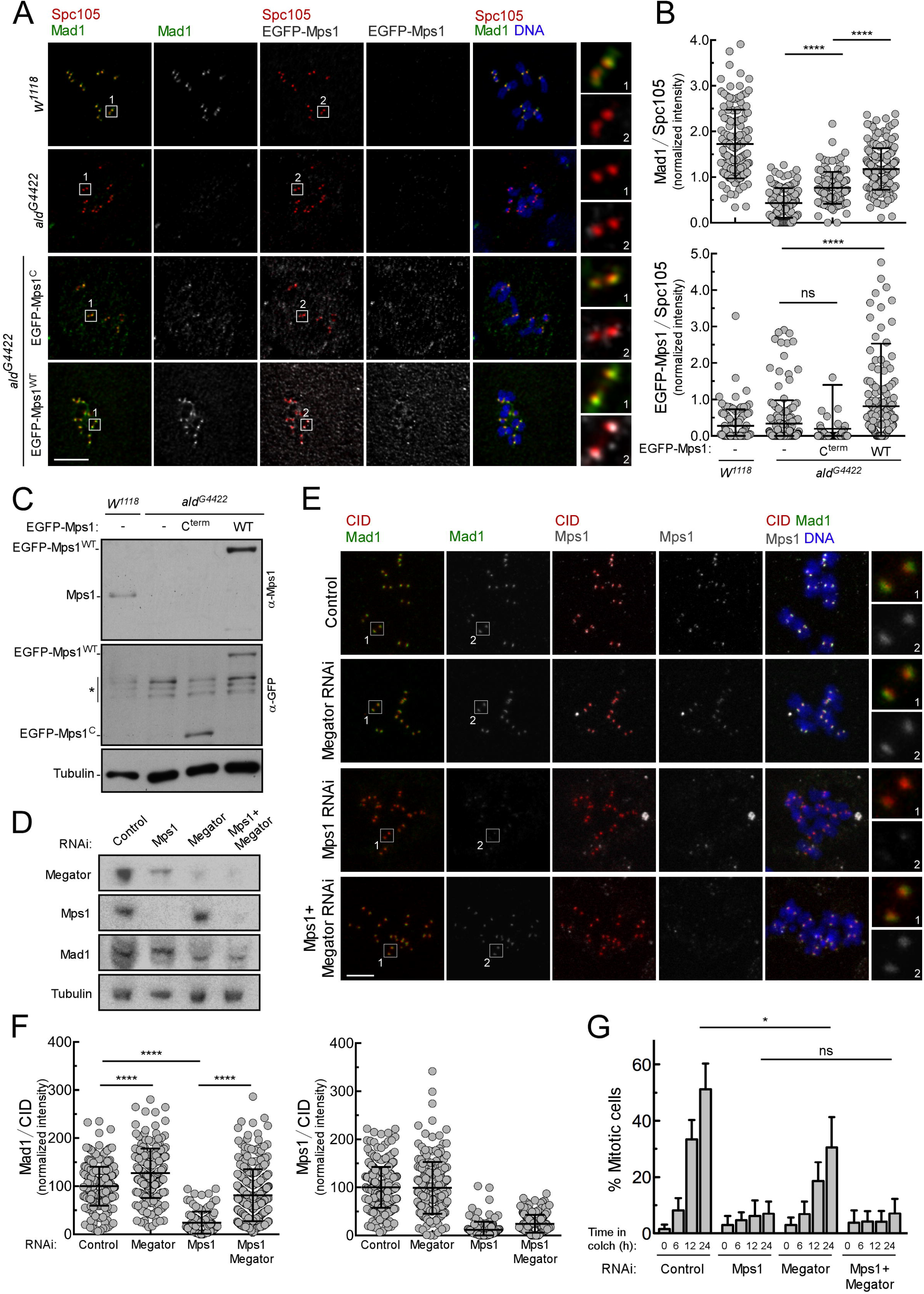
Kinetochore-extrinsic activity of Mps1 contributes for Mad1 kinetochore recruitment. (A,B) Representative immunofluorescence images (A) and corresponding quantifications (B) of Mad1 and Mps1 levels at unattached kinetochores of neuroblasts from *w^1118^* or *ald^G4422^* flies. When indicated, EGFP-Mps1-C^term^ or EGFP-Mps1-WT transgenes were expressed under control of *Mps1* native promoter in an *ald^G4422^* background. To generate unattached kinetochores, neuroblasts were incubated with colchicine (50μM) for 1.5 hours. The insets display magnifications of the outlined regions. Mad1 and Mps1 fluorescence intensities were determined relative to Spc105 signal (N ≥ 106 kinetochores for each condition). (C) Western blot analysis of endogenous Mps1, EGFP-Mps1-WT and EGFP-Mps1-C^term^ levels in total lysates of 3^rd^ *instar* larval brains from (A). (D) Western blot analysis of Mps1, Megator and Mad1 relative levels in control S2 cells and in cells depleted of the indicated proteins. Cells were incubated with MG123 (20μM) for 1 hour and with colchicine (30μM) for 2 hours. Asterisk denotes bands resulting from unspecific anti-GFP blotting. (E, F) Representative immunofluorescence images (E) and corresponding quantifications (F) of Mad1 and Mps1 levels at unattached kinetochores of control S2 cells and cells depleted of the indicated proteins. Cells were incubated with MG123 (20μM) for 1 hour and with colchicine (30μM) for 2 hours. The insets display magnifications of the outlined regions. Mad1 and Mps1 fluorescence intensities were determined relative to CID signal (N > 109 kinetochore for Mad1, N> 139 kinetochores for Mps1). (G) Mitotic index quantification based on H3^Ser10Ph^ staining of control S2 cells and cells depleted of the indicated proteins. Cells were incubated with colchicine (30μM) for time periods indicated. Data information: in (B), (F), and (G) data are presented as mean ± SD. Asterisks indicate that differences between mean ranks are statistically significant, *p<0.05, ****p<0.0001 (Kruskal-Wallis, Dunn’s multiple comparison test). Scale bars: 5μm.

**Figure S3:**
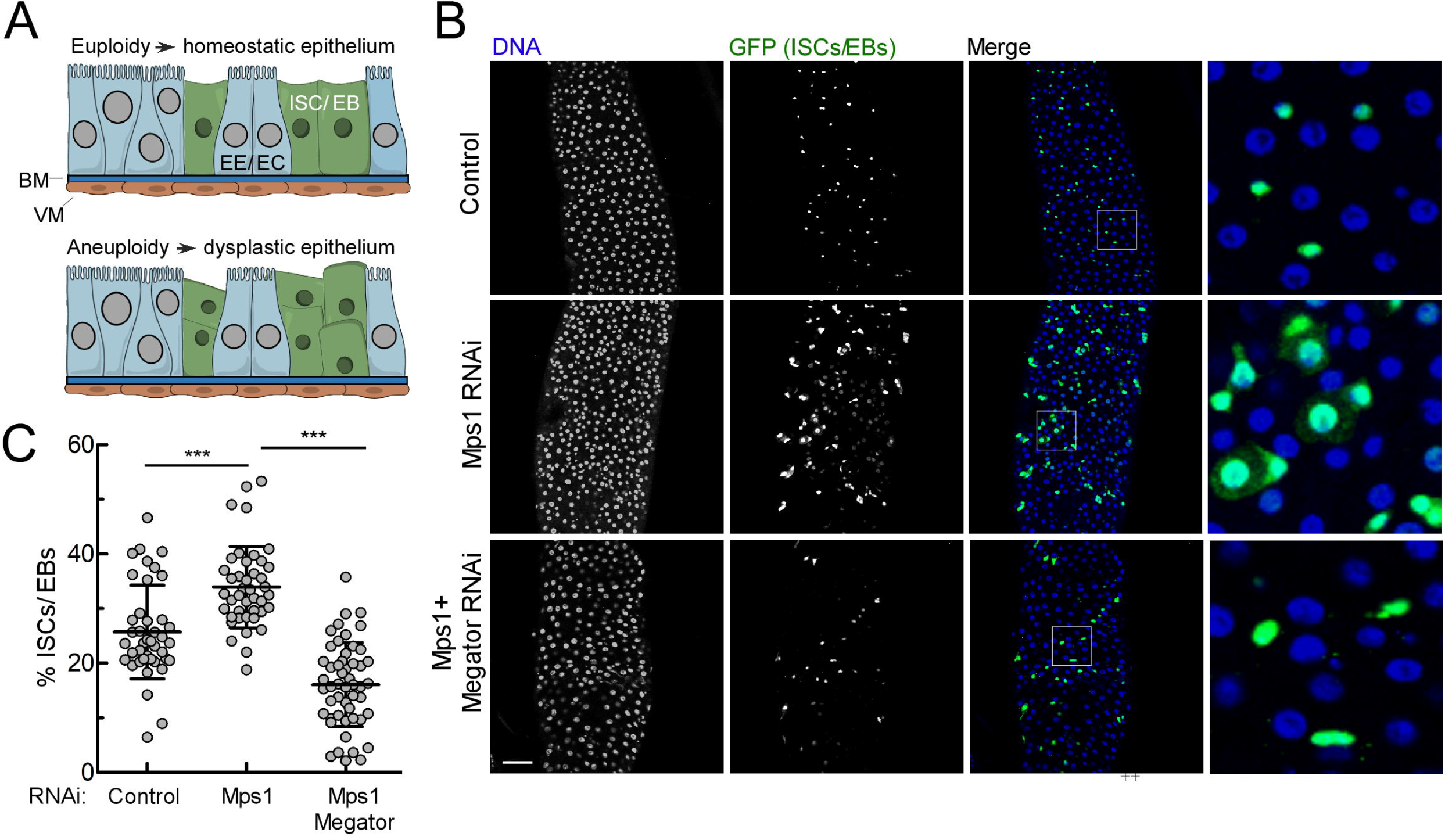
Depletion of Megator prevents intestinal dysplasia caused by lack of Mps1 activity in intestinal stem cells. (A) Schematic representation of the *Drosophila* posterior midgut epithelium under homeostatic conditions or after aneuploidy-induced dysplasia. Aneuploid ISCs/EBs over-proliferate and accumulate causing epithelium dysplasia (Resende et al., 2018). ISCs-intestinal stem cells, EBs – enteroblasts, EE –enteroendocrine cells, EC-enterocytes BM – basement membrane, VM-visceral muscle. (B,C) Representative immunofluorescence images (B) and corresponding quantifications (C) of the percentage of ISCs/ EBs (GFP-positive) in intestines with ISCs/EBs depleted of the indicated proteins. *GFP-UAS* was expressed alone or co-expressed with *UAS-MegatorRNAi* or *UAS-Mps1RNAi* under control of the *EsgGAL4* promoter during the first 20 days of adult flies. The insets display magnifications of the outlined regions. Quantification of percentage of ISCs/EBs relative to total number of cells (N ≥ 40 intestines). Data on graph represents mean ± SD. Asterisks indicate that differences between mean ranks are statistically significant, * p<0.05; *** p<0.001 (Kruskal-Wallis, Dunn’s multiple comparison test). Scale bar: 50μm.

**Figure S4:**
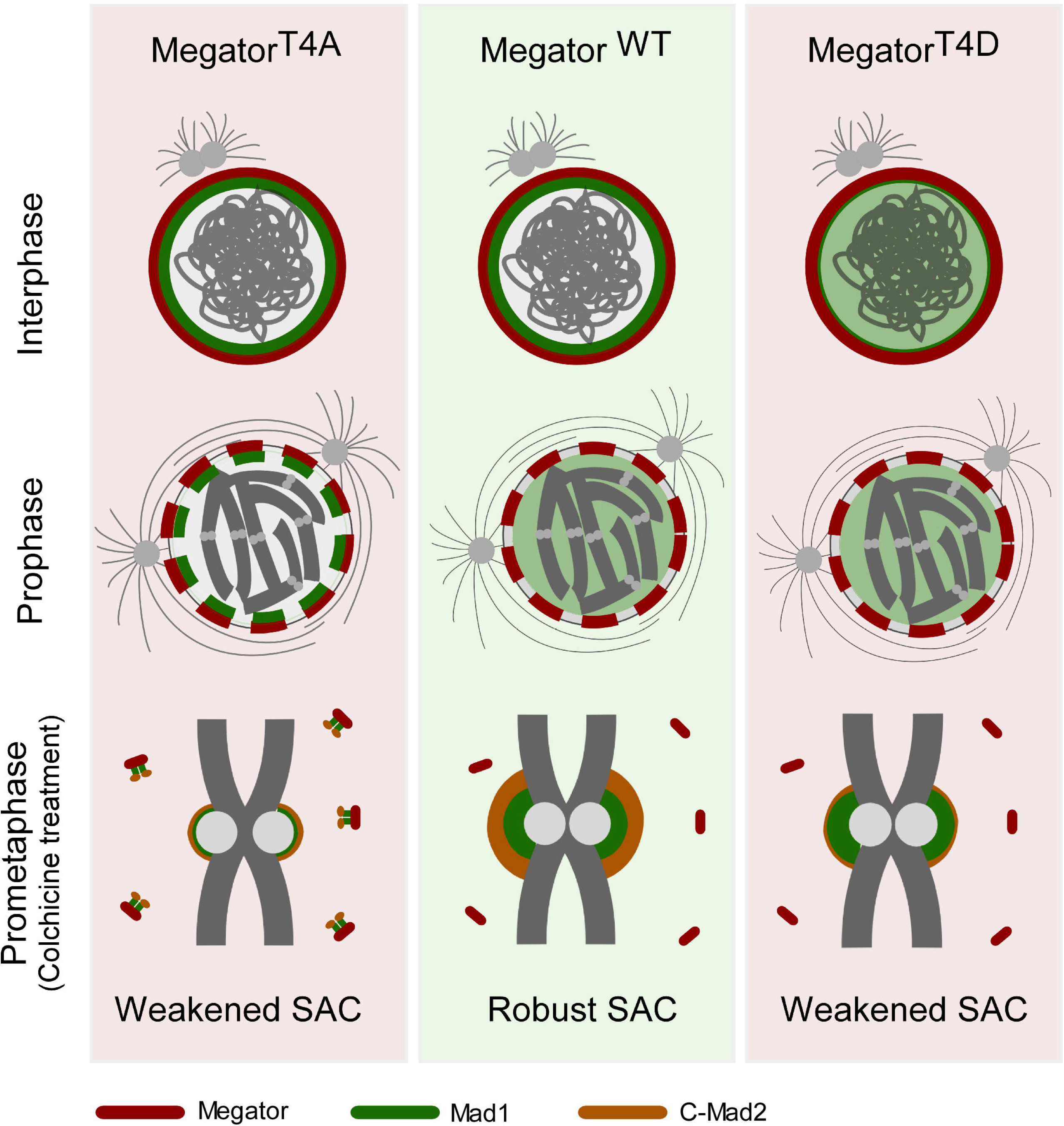
Timely phosphorylation of Megator coordinates Mad1 subcellular localization with cell cycle progression to ensure a fully-functional spindle assembly checkpoint. Preventing Mps1-mediated phosphorylation of Megator on T1259, T1295, T1338 and T1390 (Megator^T4A^) retains Mad1 associated with the nucleoporin during mitosis. This precludes proper recruitment of Mad1-C-Mad2 to prometaphase/unattached kinetochores and consequently compromises the strength of SAC signalling. On the other hand, constitutive phosphorylation of these residues (Megator^T4D^) abrogates Mad1 interaction with Megator throughout the cell cycle. Though able to efficiently accumulate Mad1 at prometaphase/unattached kinetochores, Megator^T4D^ cells also exhibit a weakened SAC function. This likely results from reduced formation of Mad1-C-Mad2 heterotetramers at NPCs during interphase, which possibly limits the assembly of pre-mitotic MCC (Rodriguez-Bravo et al., 2014) and the association of C-Mad2 with kinetochores latter in mitosis. Hence, robust SAC function requires the interaction of Mad1 with Megator at NPCs to be tightly coordinated with cell cycle progression. Activation and nuclear import of Mps1 during late G2/prophase provides a molecular switch that ensures timely release of Mad1 from NPCs precisely when kinetochores must instate SAC signalling. This kinetochore-extrinsic mechanism enables the cell to produce MCC both at NPCs in interphase and at kinetochores during mitosis, so that the checkpoint is sufficiently robust to safeguard against chromosome mis-segregation.

## References

Akera, T., Y. Goto, M. Sato, M. Yamamoto, and Y. Watanabe. 2015. Mad1 promotes chromosome congression by anchoring a kinesin motor to the kinetochore. Nat. Cell Biol. 17:1124–1133. doi:10.1038/ncb3219.

Althoff, F., R.E. Karess, and C.F. Lehner. 2012. Spindle checkpoint-independent inhibition of mitotic chromosome segregation by *Drosophila* Mps1. Mol. Biol. Cell. 23:2275–2291. doi:10.1091/mbc.e12-02-0117.

De Antoni, A., C.G. Pearson, D. Cimini, J.C. Canman, V. Sala, L. Nezi, M. Mapelli, L. Sironi, M. Faretta, and A. Musacchio. 2005. The Mad1/Mad2 complex as a template for Mad2 activation in the spindle assembly checkpoint. Curr. Biol. 15:214–225. doi: 10.1016/j.

Chen, R.H., A. Shevchenko, M. Mann, and A.W. Murray. 1998. Spindle checkpoint protein Xmad1 recruits Xmad2 to unattached kinetochores. J. Cell Biol. 143:283–295. doi:10.1083/jcb.143.2.283.

Chung, E., and R.-H. Chen. 2002. Spindle checkpoint requires Mad1-bound and Mad1-free Mad2. Mol. Biol. Cell. 13:1501–1511. doi:10.1091/mbc.02.

Collin, P., O. Nashchekina, R. Walker, and J. Pines. 2013. The spindle assembly checkpoint works like a rheostat rather than a toggle switch. Nat. Cell Biol. 15:1378–1385. doi:10.1038/ncb2855.

Conde, C., M. Osswald, J. Barbosa, T. Moutinho-Santos, D. Pinheiro, S. Guimarães, I. Matos, H. Maiato, and C.E. Sunkel. 2013. *Drosophila* Polo regulates the spindle assembly checkpoint through Mps1-dependent BubR1 phosphorylation. EMBO J. 32:1761–77. doi:10.1038/emboj.2013.109.

Cox, J., and M. Mann. 2008. MaxQuant enables high peptide identification rates, individualized p.p.b.-range mass accuracies and proteome-wide protein quantification. Nat. Biotechnol. 26:1367–72. doi:10.1038/nbt.1511.

Dick, A.E., and D.W. Gerlich. 2013. Kinetic framework of spindle assembly checkpoint signalling. Nat. Cell Biol. 15:1370–1377. doi:10.1038/ncb2842.

Emre, D., R. Terracol, A. Poncet, Z. Rahmani, and R.E. Karess. 2011. A mitotic role for Mad1 beyond the spindle checkpoint. J. Cell Sci. 124:1664–1671. doi:10.1242/jcs.081216.

Faesen, A.C., M. Thanasoula, S. Maffini, C. Breit, F. Müller, S. Van Gerwen, T. Bange, and A. Musacchio. 2017. Basis of catalytic assembly of the mitotic checkpoint complex. Nature. 542:498–502. doi:10.1038/nature21384.

Fava, L.L., M. Kaulich, E.A. Nigg, and A. Santamaria. 2011. Probing the in vivo function of Mad1:C-Mad2 in the spindle assembly checkpoint. EMBO J. 30:3322–3336. doi:10.1038/emboj.2011.239.

Hustedt, N., M. Langegger, N. Schmidt, S. Heinrich, H. Windecker, K. Sewart, and S. Hauf. 2014. Mad1 contribution to spindle assembly checkpoint signalling goes beyond presenting Mad2 at kinetochores. EMBO Rep. 15:291–298. doi:10.1002/embr.201338114.

Jelluma, N., A.B. Brenkman, I. McLeod, J.R. Yates, D.W. Cleveland, R.H. Medema, and G.J.P.L. Kops. 2008. Chromosomal instability by inefficient Mps1 auto-activation due to a weakened mitotic checkpoint and lagging chromosomes. PLoS One. 3:e2415. doi:10.1371/journal.pone.0002415.

Ji, Z., H. Gao, L. Jia, B. Li, and H. Yu. 2017. A sequential multi-target Mps1 phosphorylation cascade promotes spindle checkpoint signaling. Elife. 6:1–23. doi:10.7554/eLife.22513.

Jia, H., X. Zhang, W. Wang, Y. Bai, Y. Ling, C. Cao, R.Z. Ma, H. Zhong, X. Wang, and Q. Xu. 2015. A putative N-terminal nuclear export sequence is sufficient for Mps1 nuclear exclusion during interphase. BMC Cell Biol. 16:1–8. doi:10.1186/s12860-015-0048-6.

Kennedy, M.J., R.M. Hughes, L.A. Peteya, J.W. Schwartz, M.D. Ehlers, and C.L. Tucker. 2010. Rapid blue-light–mediated induction of protein interactions in living cells. Nat. Methods. 7:973.

Kim, D.H., J.S. Han, P. Ly, Q. Ye, M.A. McMahon, K. Myung, K.D. Corbett, and D.W. Cleveland. 2018. TRIP13 and APC15 drive mitotic exit by turnover of interphase-and unattached kinetochore-produced MCC. Nat. Commun. 9:4354. doi:10.1038/s41467-018-06774-1.

Lee, S., H. Park, T. Kyung, N.Y. Kim, S. Kim, J. Kim, and W. Do Heo. 2014. Reversible protein inactivation by optogenetic trapping in cells. Nat. Methods. 1408:363–376. doi:10.1007/978-1-4939-3512-3_25.

Lee, S.H., H. Sterling, A. Burlingame, and F. McCormick. 2008. Tpr directly binds to Mad1 and Mad2 and is important for the Mad1-Mad2-mediated mitotic spindle checkpoint. Genes Dev. 22:2926–2931. doi:10.1101/gad.1677208.

Li, C., A. Wen, B. Shen, J. Lu, Y. Huang, and Y. Chang. 2011. FastCloning: A highly simplified, purification-free, sequence-and ligation-independent PCR cloning method. BMC Biotechnol. 11:92. doi:10.1186/1472-6750-11-92.

Lince-Faria, M., S. Maffini, B. Orr, Y. Ding, C. Florindo, C.E. Sunkel, Á. Tavares, J. Johansen, K.M. Johansen, and H. Maiato. 2009. Spatiotemporal control of mitosis by the conserved spindle matrix protein megator. J. Cell Biol. 184:647–657. doi:10.1083/jcb.200811012.

London, N., and S. Biggins. 2014. Mad1 kinetochore recruitment by Mps 1-mediated phosphorylation of Bub1 signals the spindle checkpoint. Genes Dev. 28:140–52. doi:10.1101/gad.233700.113.

London, N., S. Ceto, J.A. Ranish, and S. Biggins. 2012. Phosphoregulation of Spc105 by Mps1 and PP1 regulates Bub1 localization to kinetochores. Curr. Biol. 22:900–906. doi:10.1016/j.cub.2012.03.052.

Maciejowski, J., K.A. George, M.E. Terret, C. Zhang, K.M. Shokat, and P. V. Jallepalli. 2010. Mps1 directs the assembly of Cdc20 inhibitory complexes during interphase and mitosis to control M phase timing and spindle checkpoint signaling. J. Cell Biol. 190:89–100. doi:10.1083/jcb.201001050.

Malureanu, L.A., K.B. Jeganathan, M. Hamada, L. Wasilewski, J. Davenport, and J.M. van Deursen. 2009. BubR1 N Terminus Acts as a Soluble Inhibitor of Cyclin B Degradation by APC/CCdc20 in Interphase. Dev. Cell. 16:118–131. doi:10.1016/j.devcel.2008.11.004.

Meraldi, P., V.M. Draviam, and P.K. Sorger. 2004. Timing and checkpoints in the regulation of mitotic progression. Dev. Cell. 7:45–60. doi:10.1016/j.devcel.2004.06.006.

Michalski, A., E. Damoc, J.-P. Hauschild, O. Lange, A. Wieghaus, A. Makarov, N. Nagaraj, J. Cox, M. Mann, and S. Horning. 2011. Mass spectrometry-based proteomics using Q Exactive, a high-performance benchtop quadrupole Orbitrap mass spectrometer. Mol. Cell. Proteomics. 10:M111.011015. doi:10.1074/mcp.M111.011015.

Mora-Santos, M. del M., A. Hervas-Aguilar, K. Sewart, T.C. Lancaster, J.C. Meadows, and J.B.A. Millar. 2016. Bub3-Bub1 Binding to Spc7/KNL1 Toggles the Spindle Checkpoint Switch by Licensing the Interaction of Bub1 with Mad1-Mad2. Curr. Biol. 26:2642–2650. doi:10.1016/j.cub.2016.07.040.

Moura, M., M. Osswald, N. Leça, J. Barbosa, A.J. Pereira, H. Maiato, C.E. Sunkel, and C. Conde. 2017. Protein phosphatase 1 inactivates Mps1 to ensure efficient spindle assembly checkpoint silencing. Elife. 6:1–29. doi:10.7554/eLife.25366.

Olsen, J. V., B. Macek, O. Lange, A. Makarov, S. Horning, and M. Mann. 2007. Higher-energy C-trap dissociation for peptide modification analysis. Nat. Methods. 4:709–712. doi:10.1038/nmeth1060.

Osswald, M., A.F. Santos, and E. Morais-de-Sá. 2019. Light-Induced Protein Clustering for Optogenetic Interference and Protein Interaction Analysis in Drosophila S2 Cells. Biomolecules. 9:61. doi:10.3390/biom9020061.

Primorac, I., J.R. Weir, E. Chiroli, F. Gross, I. Hoffmann, S. van Gerwen, A. Ciliberto, and A. Musacchio. 2013. Bub3 reads phosphorylated MELT repeats to promote spindle assembly checkpoint signaling. Elife. 2013:1–20. doi:10.7554/eLife.01030.

Qi, H., U. Rath, D. Wang, Y.-Z. Xu, Y. Ding, W. Zhang, M.J. Blacketer, M.R. Paddy, J. Girton, J. Johansen, and K.M. Johansen. 2004. Megator, an essential coiled-coil protein that localizes to the putative spindle matrix during mitosis in *Drosophila*. Mol. Biol. Cell. 15:4854–4865. doi:10.1091/mbc.E04.

Qian, J., M.A. García-Gimeno, M. Beullens, M.G. Manzione, G. Van der Hoeven, J.C. Igual, M. Heredia, P. Sanz, L. Gelens, and M. Bollen. 2017. An attachment-independent biochemical timer of the spindle assembly checkpoint. Mol. Cell. 68:715–730. doi:10.1016/j.molcel.2017.10.011.

Rappsilber, J., M. Mann, and Y. Ishihama. 2007. Protocol for micro-purification, enrichment, pre-fractionation and storage of peptides for proteomics using StageTips. Nat. Protoc. 2:1896–1906. doi:10.1038/nprot.2007.261.

Resende, L.P., A. Monteiro, R. Brás, T. Lopes, and C.E. Sunkel. 2018. Aneuploidy in intestinal stem cells promotes gut dysplasia in *Drosophila*. J. Cell Biol. 217:3930–3946. doi:10.1083/jcb.201804205.

Rodriguez-Bravo, V., J. Maciejowski, J. Corona, H. kon K. Buch, P. Collin, M.T. Kanemaki, J. V Shah, and P. V Jallepalli. 2014. Nuclear pores protect genome integrity by assembling a premitotic and Mad1-dependent anaphase inhibitor. Cell. 156:1017–1031. doi:10.1016/j.cell.2014.01.010.

Rodriguez-Rodriguez, J.A., C. Lewis, K.L. McKinley, V. Sikirzhytski, J. Corona, J. Maciejowski, A. Khodjakov, I.M. Cheeseman, and P. V. Jallepalli. 2018. Distinct roles of RZZ and Bub1-KNL1 in mitotic checkpoint signaling and kinetochore expansion. Curr. Biol. 28:3422–3429.e5. doi:10.1016/j.cub.2018.10.006.

Schittenhelm, R.B., R. Chaleckis, and C.F. Lehner. 2009. Intrakinetochore localization and essential functional domains of *Drosophila* Spc105. EMBO J. 28:2374–2386. doi:10.1038/emboj.2009.188.

Schweizer, N., C. Ferrás, D.M. Kern, E. Logarinho, I.M. Cheeseman, and H. Maiato. 2013. Spindle assembly checkpoint robustness requires Tpr-mediated regulation of Mad1/Mad2 proteostasis. J. Cell Biol. 203:883–893. doi:10.1083/jcb.201309076.

Scott, R.J., C.P. Lusk, D.J. Dilworth, J.D. Aitchison, and R.W. Wozniak. 2005. Interactions between Mad1p and the nuclear transport machinery in the east *Saccharomyces cerevisiae*. Mol. Biol. Cell. 16:4362–4374. doi:10.1091/mbc.E05.

Shepperd, L.A., J.C. Meadows, A.M. Sochaj, T.C. Lancaster, J. Zou, G.J. Buttrick, J. Rappsilber, K.G. Hardwick, and J.B.A. Millar. 2012. Phosphodependent recruitment of Bub1 and Bub3 to Spc7/KNL1 by Mph1 kinase maintains the spindle checkpoint. Curr. Biol. 22:891–899. doi:10.1016/j.cub.2012.03.051.

Simonetta, M., R. Manzoni, R. Mosca, M. Mapelli, L. Massimiliano, M. Vink, B. Novak, A. Musacchio, and A. Ciliberto. 2009. The influence of catalysis on Mad2 activation dynamics. PLoS Biol. 7. doi:10.1371/journal.pbio.1000010.

Sironi, L., M. Mapelli, S. Knapp, A. DeAntoni, K.-T. Jeang, and A. Musacchio. 2002. The Mad1-Mad2 complex: implications of a “safety belt” binding mechanism for the spindle checkpoint. embo J. 21:2496–2506. doi:10.1107/s0108767302093947.

Souza, C.P. De, S.B. Hashmi, T. Nayak, B. Oakley, and S.A. Osmani. 2009. Mlp1 acts as a mitotic scaffold to spatially regulate spindle assembly checkpoint proteins in *Aspergillus nidulans*. Mol. Biol. Cell. 20:2146–2159. doi:10.1091/mbc.E08.

Sudakin, V., G.K.T. Chan, and T.J. Yen. 2001. Checkpoint inhibition of the APC/C in HeLa cells is mediated by a complex of BUBR1, BUB3, CDC20, and MAD2. J. Cell Biol. 154:925–936. doi:10.1083/jcb.200102093.

Vink, M., M. Simonetta, P. Transidico, K. Ferrari, M. Mapelli, A. De Antoni, L. Massimiliano, A. Ciliberto, M. Faretta, E.D. Salmon, and A. Musacchio. 2006. In vitro FRAP identifies the minimal requirements for Mad2 kinetochore dynamics. Curr. Biol. 16:755–766. doi:10.1016/j.cub.2006.03.057.

Vleugel, M., M. Omerzu, V. Groenewold, M.A. Hadders, S.M.A. Lens, and G.J.P.L. Kops. 2015. Sequential multisite phospho-regulation of KNL1-BUB3 interfaces at mitotic kinetochores. Mol. Cell. 57:824–835. doi:10.1016/j.molcel.2014.12.036.

Yamagishi, Y., C.-H. Yang, Y. Tanno, and Y. Watanabe. 2012. MPS1/Mph1 phosphorylates the kinetochore protein KNL1/Spc7 to recruit SAC components. Nat. Cell Biol. 14:746–752. doi:10.1038/ncb2515.

Zhang, G., T. Kruse, B. López-Méndez, K.B. Sylvestersen, D.H. Garvanska, S. Schopper, M.L. Nielsen, and J. Nilsson. 2017. Bub1 positions Mad1 close to KNL1 MELT repeats to promote checkpoint signalling. Nat. Commun. 8:15822. doi:10.1038/ncomms15822.

Zhang, X., Q. Yin, Y. Ling, Y. Zhang, R. Ma, Q. Ma, C. Cao, H. Zhong, X. Liu, and Q. Xu. 2011. Two LXXLL motifs in the N terminus of Mps1 are required for Mps1 nuclear import during G 2 /M transition and sustained spindle checkpoint responses. Cell Cycle. 10:2742–2750. doi:10.4161/cc.10.16.15927.

